# The sensory coding of warm perception

**DOI:** 10.1101/502369

**Authors:** Ricardo Paricio-Montesinos, Frederick Schwaller, Annapoorani Udhayachandran, Jan Walcher, Roberta Evangelista, James F.A. Poulet, Gary R. Lewin

**Affiliations:** Department of Neuroscience, Max Delbrück Center for Molecular Medicine, Robert-Rössle Straße 10, D-13092 Berlin, Germany; Neuroscience Research Center and Cluster of Excellence NeuroCure, Charité-Universitätsmedizin, Charitéplatz 1, 10117 Berlin Germany.

**Author notes:** Correspondence to James F.A. Poulet or Gary R Lewin. Authors contributed equally (Ricardo Paricio-Montesinos, Frederick Schwaller).

**Keywords:** perception, sensory coding, warm, thermal transduction, nociception

## Abstract

Humans easily discriminate tiny skin temperature changes that are perceived as warming or cooling. Dedicated thermoreceptors forming distinct thermosensory channels or “labelled lines” are thought to underlie thermal perception. We show that mice have similar perceptual thresholds for forepaw warming to humans (~1 °C change) and do not mistake warming for cooling. Mice perform warm discrimination tasks without dedicated thermoreceptors, but use information carried by unmyelinated polymodal C-fibers. Deletion of the heat-sensitive transduction channels TRPM2 and TRPV1 did not impact warming perception or afferent coding of warm. However, without the cold sensitive TRPM8 channel, afferent coding of cooling was impaired and these mice cannot perceive warming or cooling. Our data is incompatible with the existence of thermospecific labelled lines, but can be reconciled by the existence of central circuits that compare and integrate the input from at least two types of polymodal afferents, hitherto thought to exclusively signal pain.

## Introduction

Since the discovery of hot and cold spots on the skin (Blix, 1882), the perception of innocuous warming or cooling has been hypothesized to be mediated by specific and separate sensory channels (Schepers and Ringkamp, 2010). Dedicated thermoreceptors have been described in primate skin and human skin that respond exclusively to small temperature changes and are either specific for cooling or warming (Campero et al., 2001; Hallin et al., 1982; LaMotte and Campbell, 1978). These dedicated thermoreceptors often show ongoing activity at room temperature which is usually inhibited by a temperature change in the opposite direction to the thermoreceptor preference; e.g. ongoing activity in a cooling receptor is inhibited by warming. Dedicated thermoreceptors normally have unmyelinated C-fiber axons (Darian-Smith et al., 1979, 1979; Susser et al., 1999; Yarnitsky and Ochoa, 1991), but cooling receptors with thinly myelinated Aδ-axons have been described (Campero and Bostock, 2010; Darian-Smith et al., 1973; Iggo, 1969; Susser et al., 1999). It is far from clear whether activity in dedicated thermoreceptors can alone account for warm and cool perception. Additionally, information about warming or cooling can also be relayed by so-called polymodal sensory afferents. For example, in humans, primates, and rodents there are many mechanosensitve unmyelinated C-fibers that can signal small temperature changes, but in contrast to dedicated thermoreceptors these fibers increase their firing monotonically as temperatures become noxious (Campero et al., 1996). The relative contribution of dedicated thermoreceptors as opposed to polymodal temperature sensitive afferents to the perception of innocuous cooling or warming has not been addressed.

Recently, it was found that mice are able to perceive low threshold thermal stimuli as assessed with goal-directed short-latency perception tasks (Milenkovic et al., 2014; Yarmolinsky et al., 2016). Mice are able to detect cooling of the skin with perceptual thresholds of just 1°C, very similar to those found in humans (Frenzel et al., 2012; Milenkovic et al., 2014; Stevens and Choo, 1998). We found that activity in polymodal C-fibers was required to perceive innocuous skin cooling (Milenkovic et al., 2014). It is clear that thermosensitive TRP channels activated by cooling or warming are key players in conferring temperature sensitivity to polymodal nociceptors (Caterina et al., 1997; Vandewauw et al., 2018). The availability of gene-modified mice in which specific *Trp* genes have been deleted allows the experimental manipulation of afferent temperature sensitivity. Thus, the mouse offers an ideal model to identify the precise nature of the sensory information that is necessary and sufficient for temperature perception. At the molecular level, there is overwhelming evidence that the cold activated ion channel TRPM8 is necessary for the transduction of cold by nociceptors (McKemy, 2013; McKemy et al., 2002); mice lacking this channel have severe noxious and innocuous cool-evoked behavioural and perceptual deficits (Bautista et al., 2007; Dhaka et al., 2007; Knowlton et al., 2013; Milenkovic et al., 2014). Much less is known about candidate transduction molecules for warm transduction, but recently two candidates have been identified. First, the TRPM2 channel has been shown to activated by warm temperatures (>35°C) and has been implicated as a warm transducer in sensory neurons (Tan and McNaughton, 2016; Togashi et al., 2006). Second, the capsaicin and noxious heat activated ion channel TRPV1, which is co-expressed with TRPM2 in sensory neurons (Tan and McNaughton, 2016), has been implicated in warm sensation (Song et al., 2016; Tan and McNaughton, 2016; Yarmolinsky et al., 2016). However, the expression patterns of thermosensitive TRP channels are complex in the DRG and it is clear that ion channels with opposite thermal preference (hot and cold) are co-expressed in single cells (Takashima et al., 2007; Vandewauw et al., 2018). The complexity of peripheral afferent coding of temperature prompted us to ask whether patterned sensory input or labelled lines for temperature drive distinct warm and cool perception.

Here we trained mice to report thermal stimulation of the glabrous skin of the forepaw and show that mice learn to report forepaw skin warming. In the mouse, perceptual thresholds for warm detection (~1 °C) were remarkably similar to those found in humans. By measuring sensory responses in single sensory afferent neurons in the forepaw, we could show that polymodal C-fibers activated by non-noxious warming were likely the only fibers providing information for the perceptual task. Information about cooling was also carried by overlapping as well as distinct populations of polymodal C-fibers. Genetic deletion of thermosensitive TRPs revealed that reduced cold transduction reduces or ablates warm perception, suggesting that warm sensation depends on intact cool processing. Our data are not consistent with labelled lines for warm or cold perception and reveal that distinct and specific patterns of afferent activity in polymodal nociceptors is sufficient to drive warm or cool perception.

## Results

### Warming perception in mice

To investigate the ability of mice to detect non-noxious thermal stimulation of the forepaw, we used a goal-directed thermal perception task for head-restrained mice (Milenkovic et al., 2014). A Peltier element was positioned to make constant contact with the glabrous skin of the right forepaw of water-restricted mice (Figure 1A). The Peltier device was held at a baseline temperature of 32°C and brief warming stimuli of 10°C (total duration 4s) were applied at random time intervals (Figure 1B). Mice were rewarded with a water droplet if they licked from a sensor at any time between stimulus onset to the recooling phase of the warm stimulus. If mice licked during a 2s window before the stimulus onset, a 3-30s delay was imposed as a timeout to promote stimulus-lick association. To assess whether licking was selective to the thermal stimulus, “catch” trials of the same length were used where no warming stimulus or water reward were delivered. We then compared the hit with the false alarm rates to assess performance in the training task (Figure 1B). First, we used a small Peltier element (3×3mm) to stimulate the center of the right forepaw. Mice successfully learned to report cooling of this small skin area within 3-4 days (Figure 3A) (Milenkovic et al., 2014). However, mice confronted with a warming stimulus of the same area exhibited similar hit and false alarm rates, even after 14 days of training (Figure 1C). In contrast, when we stimulated a larger skin area (Peltier area 8×8mm, covering most of the forepaw glabrous skin) mice showed high reliability in reporting warming (Figure 1D, S1A). Therefore, as in humans (Stevens et al., 1974), spatial summation of afferent input from the skin is an important factor influencing warmth perception in mice.

Next, we measured perceptual thresholds for warming by reducing stimulus amplitude after mice had learned to report a 10°C warming stimulus. We found that mice could report a temperature change of just 1°C (from 32 to 33°C; Figure 1F). These data indicate that mice can perceive tiny changes in forepaw skin temperature with similar perceptual thresholds to forearm stimulation in humans (Frenzel et al., 2012; Stevens and Choo, 1998).

**Figure 1.**
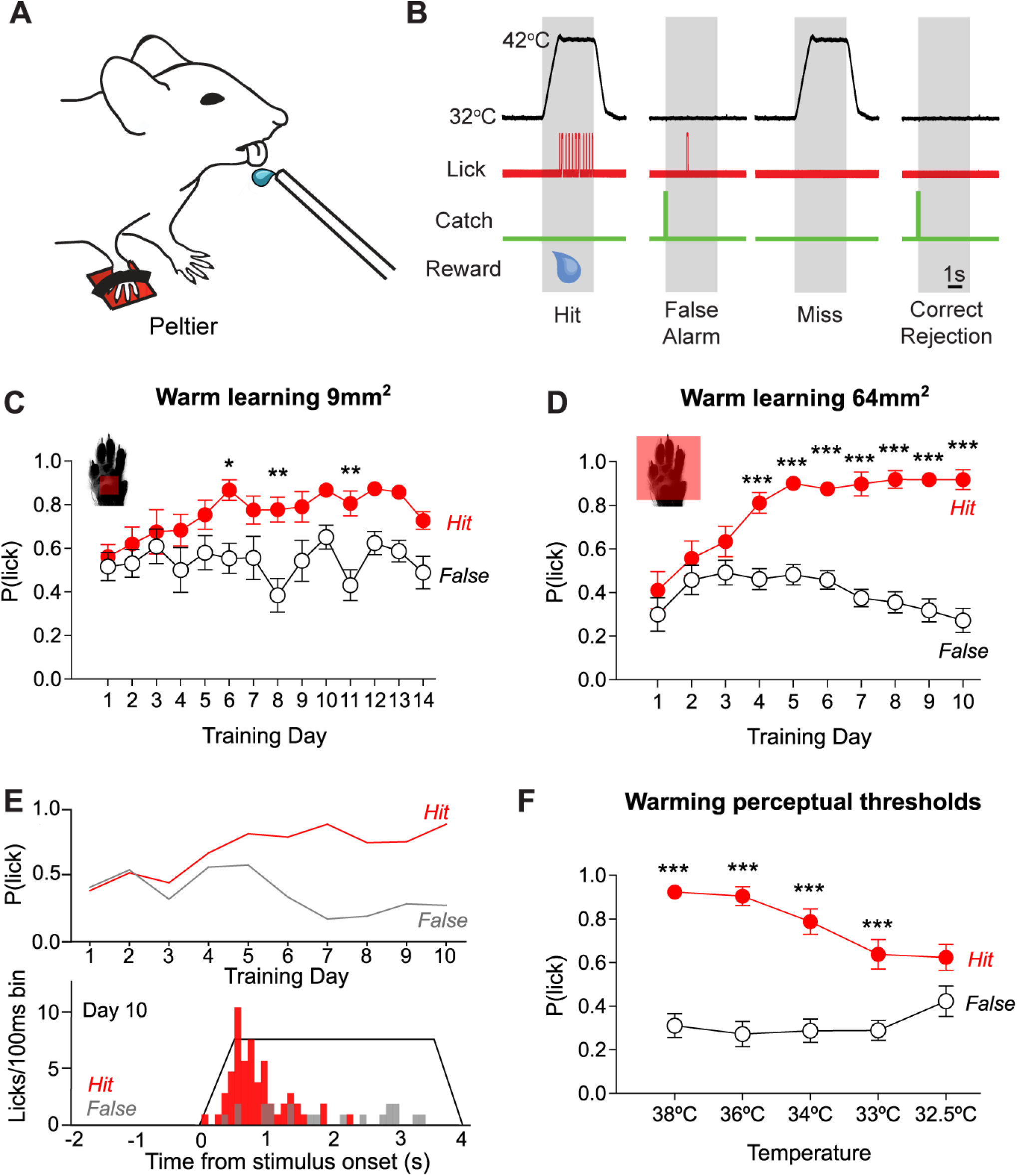
Mice learn to report non-noxious warming stimuli delivered to the forepaw. (A) Head-fixed mice report warming stimuli delivered to their right forepaw, by licking from a sensor. (B) Scheme of the warming detection task. Temperature was kept at a baseline of 32°C, and reached 42°C during warming stimuli that lasted 4 seconds. If the mouse licked within a 3.5 seconds window after onset of the stimulus, a water reward was given (Hit). We introduced “catch” trials in the same proportion, where no warming stimulus was delivered, and used this to measure spontaneous licking (False alarms). Performance was assessed by comparing the hit and false alarm rates. (C) Learning curve of warming-trained mice, using a small (3×3 mm) Peltier element shows a poor performance after two weeks of training (n=7; two-way repeated measures ANOVA with Bonferroni post-hoc tests). (D) Increasing the stimulated area with a larger (8×8 mm) Peltier improved learning performance, as mice learnt to report warming after the 4th training day (n=12; two-way repeated measures ANOVA with Bonferroni post-hoc tests). (E) Representative learning curve (top) and lick latency distribution at training day 10 (bottom) of a warming-trained mouse using a large Peltier element. (F) Decreasing stimulus amplitude over consecutive training sessions revealed a perceptual threshold of 1°C (n=12; two-way repeated measures ANOVA with Bonferroni post-hoc tests). *P < 0.05, **P < 0.01, ***P < 0.001. Data = mean ± SEM.

### Mice report forepaw warming with lower fidelity than cooling

Because mice require a larger warming stimulus area than for cooling to learn the perceptual task, we hypothesised that there may be reduced sensitivity to warming compared to cooling. To test this, we trained mice to report cooling stimuli with the large (8×8 mm) Peltier element, and found that mice immediately learned the task in the first training session (Figure 2A,B). We went on to test the cooling perceptual threshold and found that mice are able to report a cooling of 0.5°C (Figure 2C). Thus, as in humans (Stevens and Choo, 1998), cooling perception is more acute than warming.

Next, we investigated whether warm and cool stimuli have different perceptual latencies. In warming- and cooling-trained mice, peri-stimulus time histograms (PSTH) of the lick latencies illustrated that lick responses to cooling phase locked to stimulus and occurred within the first second of stimulation; however, lick responses to warming were much more variable in latency (Figure 2D-F). Comparing the development of the mean lick response latencies for warming and cooling stimuli over successive trial days, we found that warm stimuli were reported with significantly longer latencies compared to cooling throughout all training sessions (Figure 2G,H). To compare the performance of mice in our warming and cooling detection task, we used D’ (sensitivity index, see methods) measurements and found that cooling-trained mice performed better than warming-trained mice throughout all training sessions (Figure 2I). Overall, these results indicate that mice sense warming with poorer spatial and temporal resolution than cooling.

**Figure 2.**
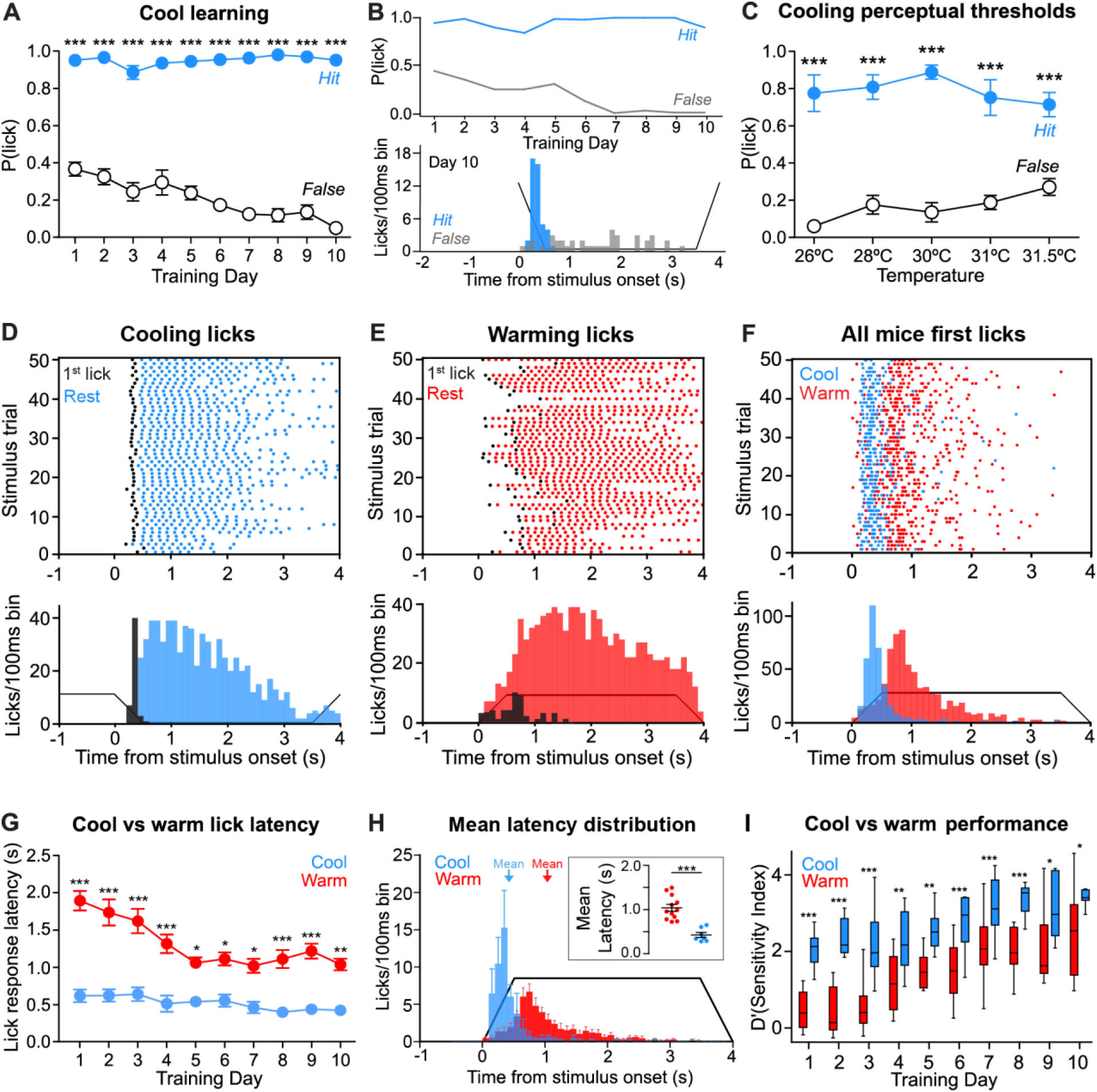
Mice report forepaw warming with lower fidelity than cooling. (A) Learning curve of cooling-trained mice revealed a high performance since the first day of training, using the large (8×8mm) Peltier element (n=7). (B) Representative learning curve (top) and lick latency distribution at training day 10 (bottom) of a cooling-trained mouse using an 8×8 mm Peltier element. (C) Perceptual threshold of cooling-trained mice was of 0.5°C (n=7, hit vs false two-way repeated measures ANOVA with Bonferroni post-hoc tests). (D) Raster plot (top) and latency histogram (bottom) of all licks made by a cooling-trained representative mouse at training day 10. The first lick of each trial is depicted in black. (E) Raster plot (top) and latency histogram (bottom) of all first licks from a warming-trained representative mouse at training day 10. (F) Raster plot (top) and latency histogram (bottom) of all first licks from all warming- and cooling-trained mice at training day 10. First licks from warming-trained mice were slower and showed a broader dispersion. (G) Mean latency of the first lick across training sessions was higher for warming- than for cooling-trained mice (warming vs cooling, n=12 and n=7 respectively; two-way repeated measures ANOVA with Bonferroni post-hoc tests). (H) Average peristimulus time histogram (PSTH) of lick latencies from warming- and cooling-trained mice at training session 10. Inset shows that mean response latency was longer for warming- (red) than cooling-trained mice (blue) at session 10 (p=0.0015, Mann Whitney U’s test, n=12 warming-trained mice, n=7 cooling-trained mice). (I) Sensitivity index (D’) measurements over training days revealed a better performance for cooling- than for warming-trained mice (warming vs cooling, n=12 and n=7 respectively; two-way repeated measures ANOVA with Bonferroni post-hoc tests). *P < 0.05, **P < 0.01, ***P < 0.001. Data = mean ± SEM. In I, boxes show median, 25% and 75% percentiles, and whiskers show minimum and maximum values.

### Mice discriminate between non-noxious warming and cooling

To investigate whether mice can discriminate warming from cooling stimuli, we inserted randomly timed cooling stimuli into a warm stimulus detection session in warming-trained mice (Figure 3A). Warming-trained mice did not report cooling stimuli; indicating that mice can correctly discriminate cooling from warming. Interestingly, warm-trained mice licked during the warming phase of the inserted cooling stimulus (Figure 3A,B). Similarly, we inserted warming stimuli into cooling detection sessions in cooling-trained mice (Figure 3C). Cooling-trained mice withheld licking to the inserted warm stimulus and only responded during the cooling phases of the warm stimulus (Figure 3D). These data indicated that, in this task, mice learnt to report a change in temperature (warming or cooling) rather than an absolute temperature value.

**Figure 3.**
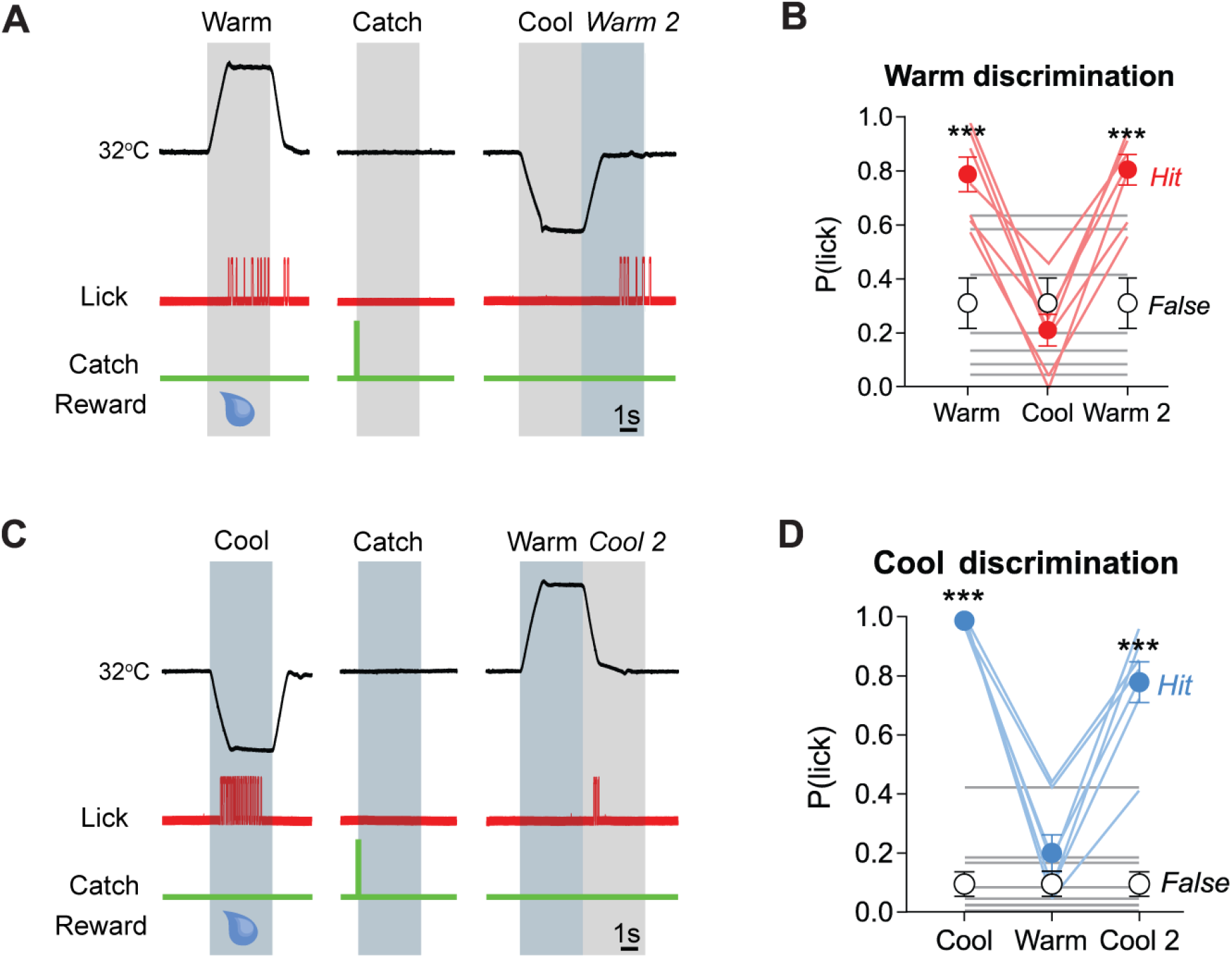
Mice discriminate non-noxious warming from cooling. (A) Scheme of the thermal discrimination task for warming-trained mice. Cool trials were introduced, and no reward was given if mice licked during cooling stimuli. Licks were also assessed during the warming phase of the right after the cooling stimulus (“Warm 2”). (B) Warming-trained mice licked the sensor during both warming types, but not during cooling stimuli (n= 7; hit vs false; two-way repeated measures ANOVA with Bonferroni post-hoc tests). (C) Scheme of the thermal discrimination task for cooling-trained mice. Warm trials were introduced, and no reward was given if mice licked during warming stimuli. Licks were also assessed during the cooling phase right after the cooling stimulus (“Cool 2”). (D) Cooling-trained mice could correctly discriminate cooling from warming, and reported cooling regardless of the absolute temperature (n=7; hit vs false two-way repeated measures ANOVA with Bonferroni post-hoc tests). *P < 0.05, **P < 0.01, ***P < 0.001. Data = mean ± SEM.

### Polymodal C-fibers detect and encode non-noxious skin warming and cooling

Next, we investigated which populations of cutaneous sensory neurons in the forepaw glabrous skin convey non-noxious warming information to the brain. We used an *ex vivo* skin-nerve preparation of the medial and ulnar nerves innervating the glabrous skin of the forepaw (Walcher et al., 2018) and examined which afferent fibers detect skin warming stimuli in a range relevant for the perceptual performance of the mouse (Figure 4A). Thermal stimulation of forepaw single-units was achieved using a Peltier device with a continuous 1°C/s warming ramp from 32-48°C and a cooling ramp of 1°C/s from 32-12°C. Once thermally activated units were identified and characterized, they were further classified based on their axonal conduction velocity and response to other modalities (mechanical and cooling stimuli). We found that the vast majority of thermo-sensitive afferents in the mouse forepaw glabrous skin (33/35) had conduction velocities below 1 m/s and were therefore classified as C-fibers (Figure S2A). The remaining two thermosensitive afferents had conduction velocities in the Aδ-fiber range (1-10 m/s), and one was classified as a mechanoheat-sensitive afferent (A-MH) and the other as mechanocold (A-MC) (Figure S2A). Thermosensitive C-fibers were almost all polymodal and classified according to the response properties as C-mechanoheat (C-MH; 19/35), C-mechanoheatcold (C-MHC; 6/35), or C-mechanocold (C-MC; 6/35) fibers. Two C-fibers were found to respond only to cold, so-called C-cold fibers (C-C), these fibers did not have the physiological properties of dedicated thermoreceptors as they had no ongoing activity and high thresholds (Figure S2A). Notably, at rest, thermosensitive-fibers were not spontaneously active when the skin temperature was held at 32°C. Some C-fibers would display spontaneous activity at rest after multiple exposure to high intensity heat stimuli (48°C; data not shown). During a 1°C/s heating ramp from 32-48°C, there was spike activity during the (non-noxious) warm phase of the ramp in a sub-population of C-MH and C-MHC afferents (Figure 4B,C). Notably, all fibers increased their firing rate as temperature increased into the noxious range, demonstrating that warm-sensitive afferents are not preferentially activated by non-noxious warming (Figure 4B, D). Heat-responsive afferents had a broad spectrum of heat-thresholds, ranging between 33-47°C, but activity during the warm phase of the ramp was sparse (Figure 4C,D).

Next, we stimulated the receptive fields of thermoreceptors with a series of 4s warming and cooling ramps with the same temperature increments and time course used for behavioural experiments (Figure 4E). Similar to a continuous heat ramp stimulus (Figure 4B,D), C-fiber spike rate increased as the temperature of the ramp increased (Figure 4F). Interestingly, the 32-42°C warming ramp used for the warm perception task evoked spike activity in 54% of all thermoreceptors (19/35) and in the majority of heat-sensitive C-fibers (22/25 heat-sensitive fibers) in the forepaw skin (Figure 4F; example shown in Figure 4E). Two warm-sensitive C-MH fibers were found to be activated by 32-33°C heat ramp (Figures 4F,G), the smallest temperature difference that can be reliably detected by the mouse (Figure 1F); each firing one action potential per stimulus (example shown in Figure 4G). Fewer cool-responsive fibers were found compared to warming (34%), and only one fiber was found to be activated at 30°C (Figure S2C,D,E). Peristimulus time histograms of spike latency during warm step (32-42°C) and cool step (32-22°C) stimuli demonstrated that warm and cool-evoked spiking had similar temporal features in warm and cool-sensitive C-MH, C-MHC and C-MC fibers (Figure 4H). These data demonstrate that skin warming and cooling is coded by overlapping populations of polymodal C-fibers. Notably, only a small subset of these afferents displayed sparse firing activity in response to <2°C change of temperature. Since the mouse needs to integrate spatial information from almost the entire forepaw glabrous skin, our data is consistent with a model where innocuous warming is detected by integrating information from large numbers of sparsely coding warm sensitive afferents.

**Figure 4.**
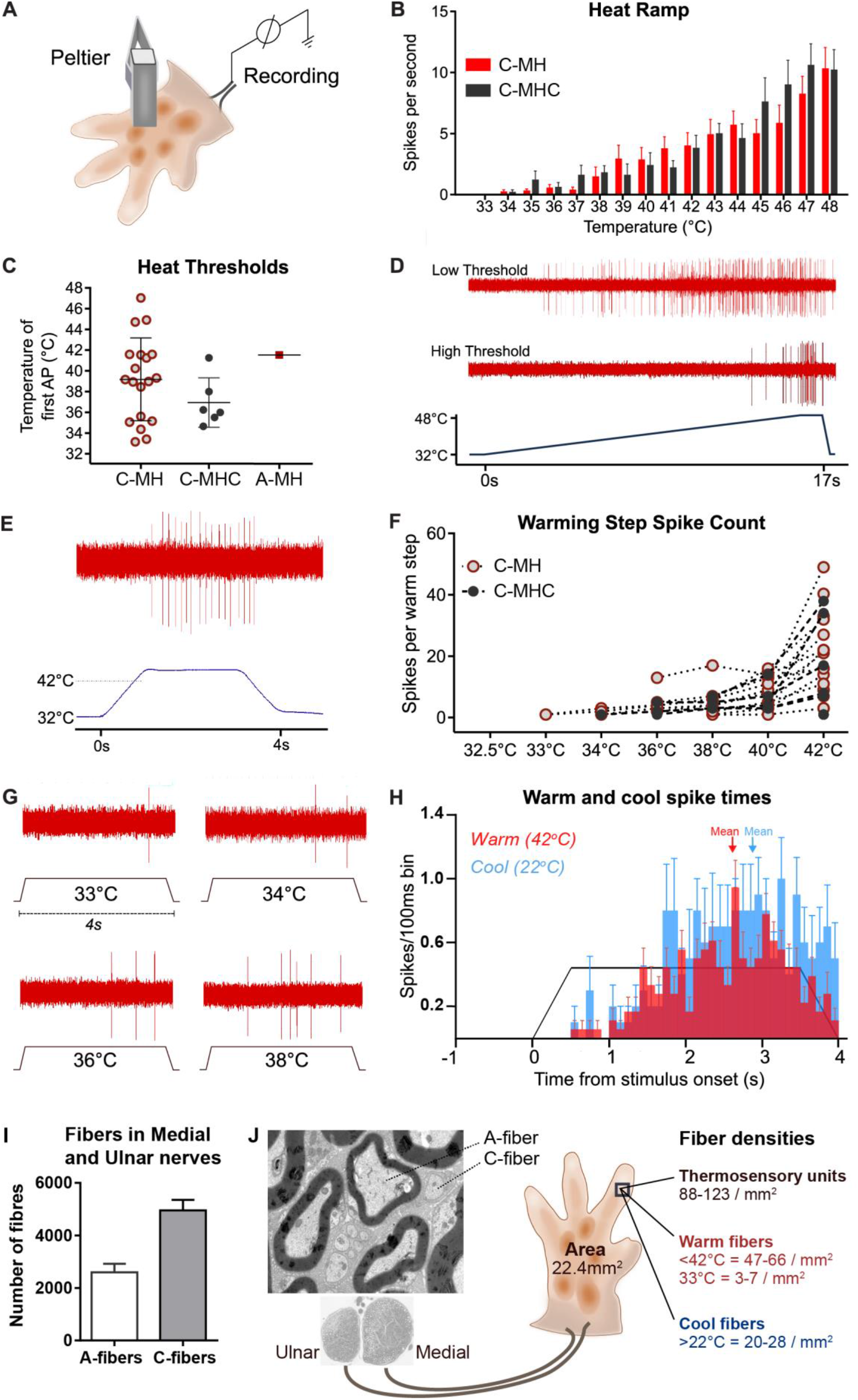
Forepaw cutaneous thermosensory neuron recordings. (A) Schematic of forepaw glabrous thermosensory cutaneous afferents recordings using the ex vivo skin-nerve preparation. N=35 units were recorded from 10 animals. (B) Mean action potential firing of populations of C-MH and C-MHC fibers during 1°C/s heat ramp. Data = mean ± SEM. (C) Heat thresholds of the different heat-sensitive fiber subtypes recorded. Data = mean ± SEM. (D) Representative responses of two different thermosensitive fibers, one low threshold and one high threshold, to temperature increase. (E) Representative afferent recording showing responses to the same warming stimulus used in the warming detection behaviour task. (F) Individual C-MH and C-MHC firing activity increased in response to warming steps of different amplitude used in behaviour threshold experiments. All warming-sensitive afferents increased firing rate as temperature increased. (G) Representative recordings showing afferent responses to warming of different amplitudes. Two C-fibers were activated by 33°C warming stimulus, each firing on average one action potential. (H) PTSHs of warm and cool-sensitive afferents responding to 32-42 °C warm stimulus (red) or 32-22°C cool stimulus (blue). (I) Mean number of A and C-fibers in the Medial and Ulnar nerves which innervate the forepaw (n=4). (J) Schematic diagram of thermosensory fiber densities in the mouse forepaw, extrapolated from the number of fibers in the Medial and Ulnar nerves (example electron micrograph shown), paw skin area, and from physiology data in figure 4.

### The density of warm sensitive fibers in the forepaw

In order to estimate the density of thermoreceptor innervation of the forepaw glabrous skin we counted the number of myelinated and unmyelinated fibers in the Medial and Ulnar nerves from transmission electron micrographs (Figure 4I,J). Assuming that the vast majority of axons in these two nerves innervate the glabrous forepaw skin we estimated the innervation density to be ~116 A-fibers/mm2 and 176 C-fibers/mm2 of skin. Extrapolating from functional data obtained from Median and Ulnar nerve recordings we estimate that there are 88-123 thermosensitive C-fibers/mm2 (50-70% of cutaneous C-fibers), 47-66 warm-sensitive fibers/mm2 and 20-28 cool-sensitive fibers/mm2 (Figure 4J). Thus we predicted that ~3-7 fibers/mm2 would be activated at 33°C, the perceptual threshold for warming (Figure 4J). Based on these density estimates a warming stimulus from 32-42°C applied to an area of 9mm2 would activate 423-594 warm fibers, but was insufficient for mice to reliably perceive warming (Figure 1C). The larger Peltier element (that stimulates a skin area of 22.4 mm2) would activate between 1044-1481 fibers with a 32-42°C warming stimulus sufficient to drive reliable warmth detection (Figure 1D).

### Normal warmth perception requires thermosensitive TRP channels

Whilst many thermally-gated ion channels have been shown to be activated by heating *in vitro* (Vriens et al., 2014), there is some debate over which channels are required for warm detection. Recent studies have provided evidence for both TRPV1 and TRPM2 involvement in warm detection (Yarmolinsky et al., 2016b) (Song et al., 2016; Tan and McNaughton, 2016). We therefore used mice with targeted null mutations in candidate TRP channels to ask which are required for sensory coding of warm and its perception. We trained mutant and wild type mice (both back-crossed onto C57bl/6 backgrounds) to detect warm stimuli to the forepaw using the larger Peltier device (8×8mm). We found that *Trpv1^−/−^* mice successfully learned to report non-painful warming stimulation of the forepaw (32-42°C) (Figure 5A). When comparing the performance of wild type and *Trpv1^−/−^* mice in the warming task using the D’ sensitivity index, we found no significant differences (Figure 5F), and no differences in lick-response latencies (Figure S1B). We next determined the mean warm detection threshold of *Trpv1^−/−^* mice by reducing the amplitude of the warming stimulus over subsequent testing days. Just like wild type mice, *Trpv1^−/−^* mice could detect a temperature change of 1°C (32-33°C; Figure 5B). Together, these findings suggest that TRPV1 is dispensable for warm perception.

*Trpm2^−/−^* mice were also able to learn to report non-painful warming stimulation of the forepaw (32-42°C) over the 10-day training period (Figure 5C). However, we found that learning performance, as measured by D’, was significantly impaired in *Trpm2^−/−^* mice with a lower sensitivity on days 8 to 10 compared to wild type control mice (Figure 5F). Additionally, there was a more pronounced spread of the mean lick latencies to 32-42°C in *Trpm2^−/−^* mice compared to wild type mice (Figure S1C). Moreover, *Trpm2^−/−^* mice had slightly higher warming perceptual thresholds (2°C) compared to wild type (1°C) (Figure 5D). These data indicate that TRPM2 has a role in non-noxious warming perception, but is not essential for mice to perceive warmth.

It is established that TRPM8 is a transducer of skin cooling which is required for cold avoidance (Bautista et al., 2007; Colburn et al., 2007; Dhaka et al., 2007) and perception (Milenkovic et al., 2014). TRPM8, however, is co-expressed with other TRP channels, such as TRPV1, in single cells (Dhaka et al., 2008; Takashima et al., 2007). Data presented here for the forepaw as well as literature from rodent hind paw clearly demonstrates that many C-fiber afferents can respond to both warming and cooling (Lewin and Mendell, 1994; Milenkovic et al., 2014). We therefore wondered whether TRPM8 could play a role in warmth perception. Surprisingly, we found that *Trpm8^−/−^* mice completely failed to learn to report forepaw warming stimuli (32-42°C) during the 10-day training task (Figure 5E). False-licking rates remained similar to hit rates over the training session, licking was poorly correlated to the stimulus time (Figure S1D) and D’ measurements were significantly reduced compared to wild type mice (Figure 5F). However, *Trpm8^−/−^* mice easily learned to report mechanical stimuli applied to the forepaw and auditory stimuli with short lick response latencies (Figure S1E-G), demonstrating that the warmth perception deficit is not due to a general impairment in learning.

**Figure 5.**
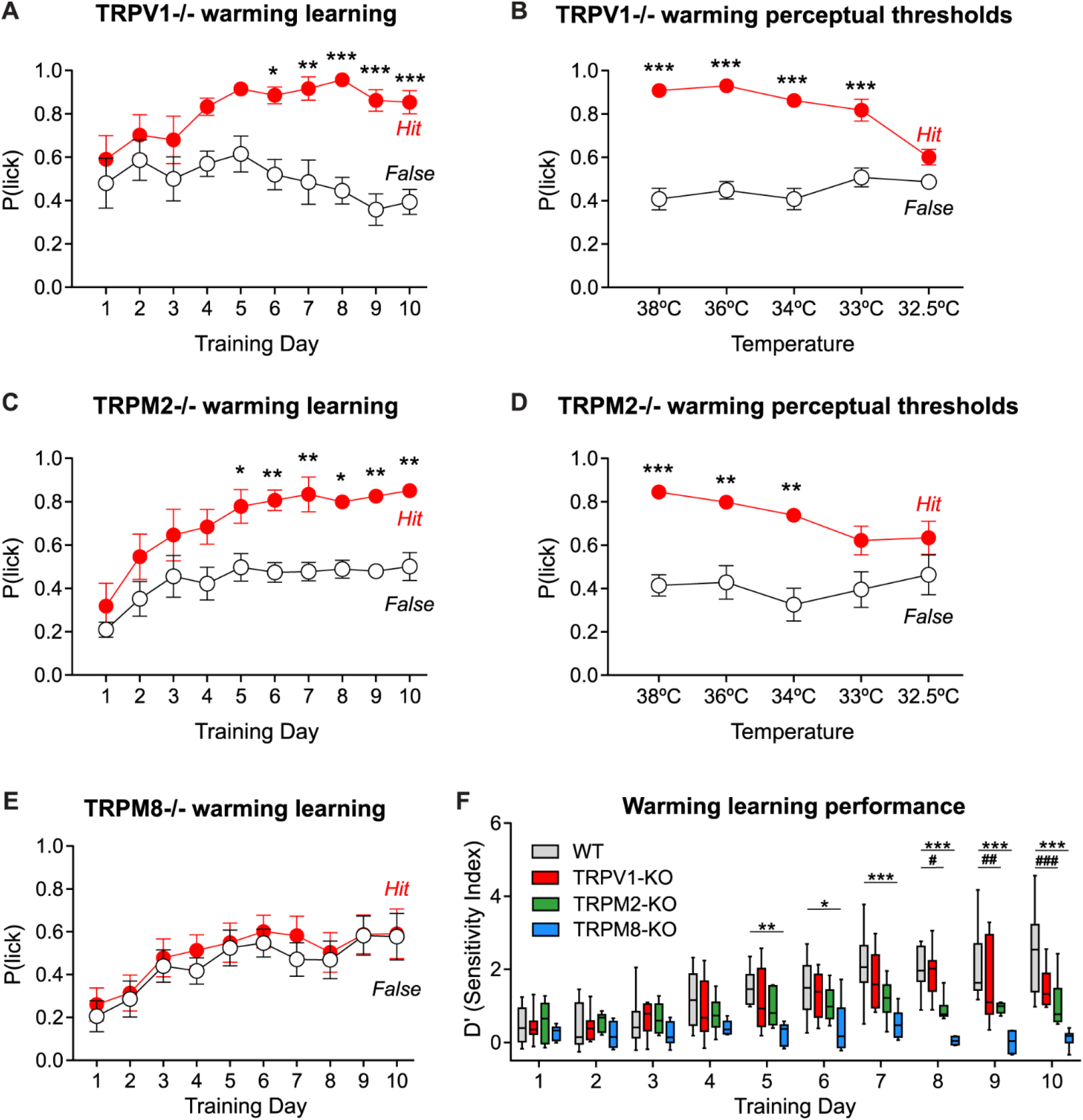
Warm behavior training in *Trpv1^−/−^*, *Trpm2^−/−^ and Trpm8^−/−^* mice. (A) *Trpv1^−/−^* mice learn to report 32-42°C forepaw warming in the behavior training task (n=8; hit vs false, two-way repeated measures ANOVA with Bonferroni post-hoc tests). (B) *Trpv1^−/−^* mice could detect warming stimuli of as little as 1 degree, similarly to wild type mice (n=8; hit vs false, two-way repeated measures ANOVA with Bonferroni post-hoc tests). (C) *Trpm2^−/−^* mice also learn to detect forepaw warming in the training task (n=6; hit vs false, two-way repeated measures ANOVA with Bonferroni post-hoc tests). (D) *Trpm2^−/−^* mice could detect warming stimuli of 2°C, but not below (n=6; hit vs false, two-way repeated measures ANOVA with Bonferroni post-hoc tests). (E) *Trpm8^−/−^* mice showed a complete absence of learning in the forepaw warming detection task, as hit and false alarm rates remained similar over the training sessions (n=10; hit vs false, two-way repeated measures ANOVA with Bonferroni post-hoc tests). (F) Sensitivity index (D’) measurements over training days revealed that *Trpm2^−/−^* mice (#) and *Trpm8^−/−^ mice* (*), but not *Trpv1^−/−^* mice, showed impaired performance in reporting forepaw warming (control vs *Trpv1^−/−^*, *Trpm2^−/−^ or Trpm8^−/−^*, two-way repeated measures ANOVA with Bonferroni post-hoc tests). ^*, #^P < 0.05, ^**, ##^P < 0.01, ^***, ###^P < 0.001. Data = mean ± SEM. In F, boxes show median, 25% and 75% percentiles, and whiskers show minimum and maximum values.

### Sensory afferent coding of warming and thermosensitive TRP channels

We next investigated whether genetic ablation of *Trpv1*, *Trpm2* or *Trpm8* alters the ability of C-fibers to detect skin warming or noxious heat. We found that C-fibers in the hairy and glabrous skin of the hind paw display responses to warming and cooling that were indistinguishable from those of the forepaw. We used the *ex vivo* skin-nerve preparation of the saphenous nerve innervating the hind paw in wild type control and *Trp* mutant mice. We recorded a total of 63 thermosensitive afferent fibers from wild type mice and 37 thermosensitive fibers from *Trpv1^−/−^* mice, again using warming ramps as the primary search stimulus. All fibers recorded were polymodal responding to combinations of mechanical stimuli and heat and/or cold. We found no afferent Aδ-fibers or C-fibers in our data set that responded only to warming or only to cold, i.e. dedicated thermoreceptors (Figure 6A). The receptive fields of thermoreceptors were stimulated using a 1°C/s heating ramp from 32–48°C and a cooling ramp from 32–12°C.

In wild type mice we found that, on average, hind paw C-MHC fibers have lower heat-thresholds compared to C-MH fibers (mean thresholds: C-MHC 39.60°C; C-MH 42.36°C) (Figure S3D), and show more spiking activity during non-noxious warming from 32-42°C (Figure S3E). Consistently, most of the C-MHC in the hairy skin (16/22) or forepaw (6/6) showed spiking during non-noxious warming. However, noxious heat and mechanical evoked firing rates were comparable between C-MH and C-MHC fibers, and not significantly different (Figure S3B,C,F). Thus, as a population, C-MHC fibers may be more responsive to skin warming than C-MH fibers.

When comparing recordings of C-fibers from wild type and *Trpv1^−/−^* mice we found no differences in the proportion of C-MH, C-MHC or C-MC fibers (Figure 6A), or in the heat thresholds of thermosensitive C-MH and C-MHC fibers (Figure 6B,C). We then investigated the thermally evoked spiking activity using our standard 32–48°C ramp heat ramp in afferents recorded from *Trpv1^−/−^* mice, and observed that spiking rates at noxious temperatures (>44°C) were significantly lower in C-MH fibers recorded from *Trpv1^−/−^* mice compared to wild type controls (Figure 6D). Interestingly, there was no deficit in noxious heat spiking rates evoked from C-MHC fibers recorded from the same *Trpv1^−/−^* mice (Figure 6E). Importantly, when we measured mean spiking rates to thermal stimuli that can be considered non-noxious (the 32-42°C used for training), we found that there was no significant difference in the mean total spike rates of both C-MH and C-MHC fibers during the non-noxious warming phase of the ramp, between *Trpv1^−/−^* and control mice (Figure 6D-H). We also quantified cool-evoked firing activity of C-MHC and C-MC fibers during the 32–22°C cooling phase of the 32–12 °C ramp: afferent responses were not altered in *Trpv1^−/−^* mice compared to controls (Figure 6J,K). We also recorded from 70 C-fibers from control hind paw glabrous skin and the data to from 82 C-fibers recorded from *Trpv1^−/−^* mice, the results broadly matched those from hairy skin (data not shown). Together, these findings show that the presence of TRPV1 channels in C-MH fibers is essential for normal encoding noxious heat, but is not necessary for the detection of non-noxious skin warming.

We next examined the thermal sensitivity of C-fibers in *Trpm2^−/−^* mice and recorded from 37 thermosensitive C-fiber units and compared their properties to the 63 control units from wild type mice. We found that the proportion of cool-sensitive fibers (C-MHC and C-MC fibers) was significantly lower in *Trpm2^−/−^* mice (Figure 6A). However, heat and cold temperature thresholds of C-MH and C-MHC fibers were not significantly different between afferents recorded from control and *Trpm2^−/−^* mice (Figure 6B,C, S4D). Spike rates of C-MH and C-MHC fibers during the 1°C/s heat ramp from 32–48°C were not different between afferents recorded from *Trpm2^−/−^* and control mice (Figure 6D,E). Similarly, quantification of the total spikes evoked from C-MH and C-MHC fibers during the 32-42°C warm phase of the ramp revealed no significant differences between *Trpm2^−/−^* and wild type afferents (Figure 6G,H). Cool-evoked firing activity of C-MHC and C-MC fibers during 32–22°C cooling was also not different between *Trpm2^−/−^* and control mice (Figure 6J,K). Overall, these data demonstrate that the presence of TRPM2 is not absolutely required for warm sensitivity of cutaneous C-fibers.

Finally, we analysed C-fiber afferents in *Trpm8^−/−^* mice and using 1°C/s 32-48°C heat and 32–12°C cold ramps. We recorded 32 thermosensitive C-fibers from *Trpm8^−/−^* mice and compared their stimulus-evoked responses to the 63 units from wild type control mice. Similar to previous findings (Bautista et al., 2007; Milenkovic et al., 2014), we found a reduction in the incidence of cold-sensitive C-MHC and C-MC fibers in *Trpm8^−/−^* mice compared to wild type (Figure 6A). Furthermore, the few remaining cold-sensitive C-MHC and C-MC fibers showed substantially reduced total firing activity during the 32–22°C cooling ramp (Figure 6I,J,K). The mean temperature thresholds for spiking to both warming and cooling were, however, not significantly different between control and *Trpm8^−/−^* C-fibers (Figure 6B,C, S4F). We quantified afferent responses during the 32–48°C heat ramp and found that C-MH and C-MHC fibers demonstrated that the mean heat-evoked firing rates were indistinguishable between fibers in control and *Trpm8^−/−^* mice (Figure 6D,E). Similarly, total mean spike numbers during the 32-42°C warming phase of the temperature ramp were not different in both C-MH and C-MHC-fibers between control and *Trpm8^−/−^* mice (Figure 6G,H). Thus despite an apparent loss of warm perception in *Trpm8^−/−^* mice, skin warming can still be detected and encoded by polymodal C-fibers in these mice.

**Figure 6.**
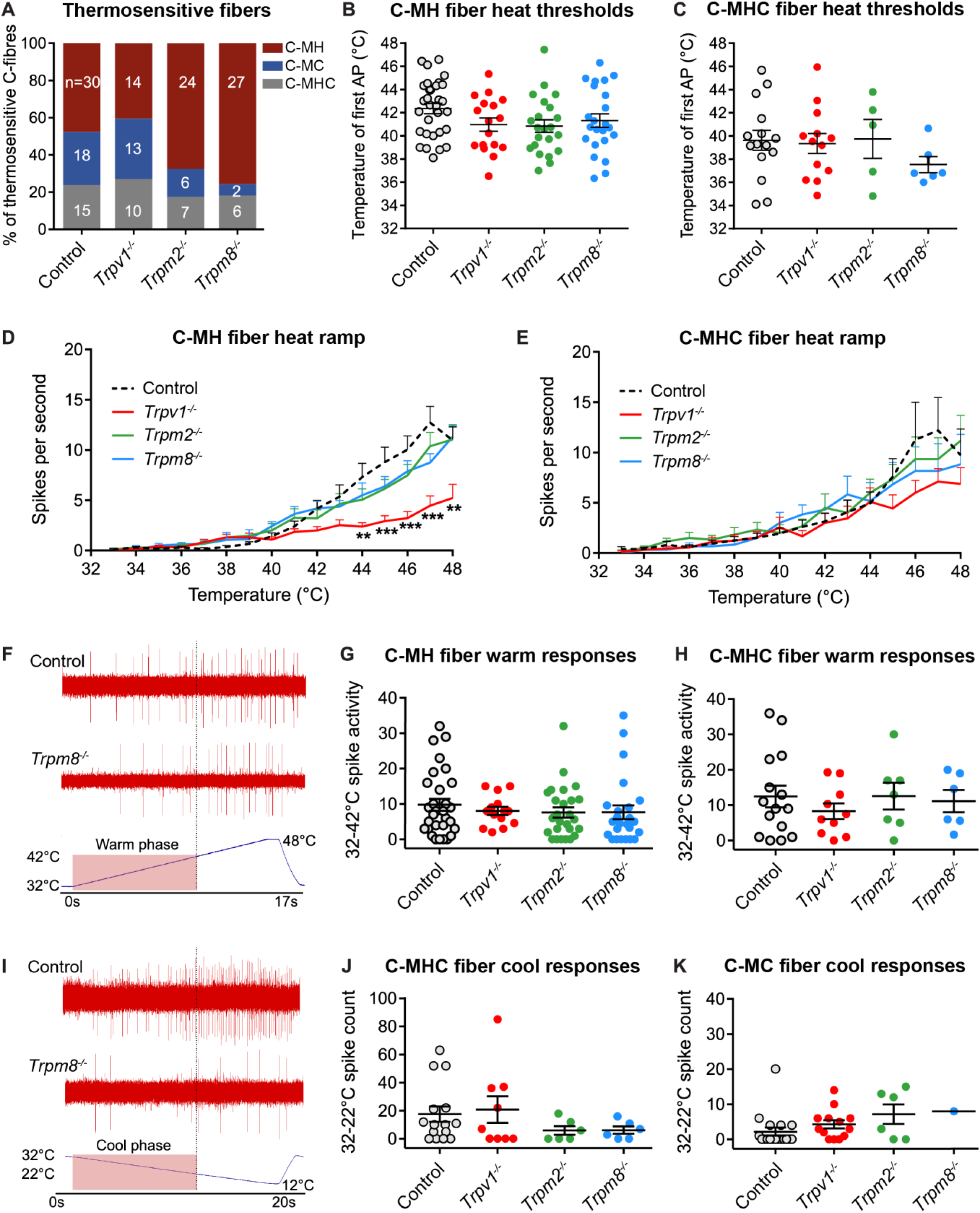
Cutaneous thermosensory neurons in control, *Trpv1^−/−^*, *Trpm2^−/−^ and Trpm8^−/−^* mice. Cutaneous thermosensory neurons were recorded in the saphenous nerve innervating the hindpaw using the ex vivo skin nerve preparation. (A) Proportion and total numbers of thermosensitive C-MH, C-MC and CMHC fiber populations in wild type control (n=63 units from 9 animals), *Trpv1^−/−^* (n=37 units from 7 animals), *Trpm2^−/−^* (n=37 units from 6 animals) and *Trpm8^−/−^* (n=32 units from 7 animals) mice. (B) Heat thresholds of C-MH fiber populations in the different mutant mouse groups were not significantly different to control (one-way ANOVA with Bonferroni post-hoc analysis). (C) Heat thresholds of CMHC fiber populations were also not different between groups. (D) During 32–48°C heat ramp, spike activity of *Trpv1^−/−^* C-MH fibers was significant lower than control at temperatures higher than 43 °C (two-way ANOVA with Bonferroni post-hoc analysis), but not for *Trpm2^−/−^* or *Trpm8^−/−^* C-MH populations. (E) C-MHC fiber spike activity during 32–48°C heat ramp was similar between groups. (F) Representative recording trace from a single control and *Trpm8^−/−^* C-MH fiber during 32–48°C heat ramp. Total spike activity in (G) and (H) was calculated during the 32-42°C warm phase of the ramp. (G) C-MH fiber population responses to the warm phase of the ramps were not different between groups. (H) Similarly, C-MHC fiber population warm responses were similar between genotypes. (I) Representative recording trace from a single control and *Trpm8^−/−^* C-MHC fiber during 32–12°C cooling ramp. Firing activity quantified during the 32–22°C cool phase is highlighted. (J) Total spike activity of C-MHC fibers during the 32–22°C cool phase of the cold ramp did not significantly differ between groups. (K) Total spike activity of *Trpv1^−/−^* and *Trpm2^−/−^* C-MC fibers during the 32–22°C cool phase of the cold ramp did not from control group. In contrast, only 1 *Trpm8^−/−^* C-MC fibers was found and had low cold-evoked spike activity compared to control. Due to low C-MC fiber numbers in the *Trpm8^−/−^* group, statistical analysis was not possible. *P < 0.05, **P < 0.01, ***P < 0.001. Data = mean ± SEM.

### Pharmacological inactivation of TRPM8 impairs warming perception

The loss of warm sensation in *Trpm8^−/−^* mice could be an indirect consequence of the early developmental loss of cooling information reaching the brain, as TRPM8-dependent cooling detection is absent throughout development. Another possibility is that functional TRPM8 channels are required to provide information about temperature changes to enable warm perception. We addressed the likelihood of these two scenarios by acutely inactivating TRPM8 in the forepaw of wild type mice, using PBMC (1-Phenylethyl(2-aminoethyl)(4-benzyloxy)-3-methoxybenzyl)carbamate), a selective antagonist of TRPM8 that has been shown to suppress cooling-responsive cells and to reduce cooling-evoked behavioural responses in mice (González et al., 2017; Knowlton et al., 2011; Yudin et al., 2016). We first trained wild type animals to report warming stimuli and, once mice were able to successfully report warming, we pharmacologically inactivated TRPM8 by performing a transdermal injection in the plantar side of the right forepaw and tested their warming perception ability (Figure 7A,B). Twenty minutes after subcutaneous PBMC injection into the forepaw, mice showed a significantly poorer warm detection performance compared to DMSO control-injected mice as shown by reduced D’ indices (Figure 7C,D). Furthermore, the latencies to report the stimuli in the successful hit trials were longer when mice were given local PBMC (Figure 7E). PSTH analysis further highlighted a warming perception deficit induced by TRPM8 blockade, as seen by the similarity of the hit and false alarm latency distributions (Figures 7F,G). Interestingly, these effects were reversible as mice showed baseline levels of performance and latencies one day after PBMC injection in the forepaw (Figures 7C-E). Together, these data suggest that functional TRPM8 channels are acutely required for the normal warm perception.

**Figure 7.**
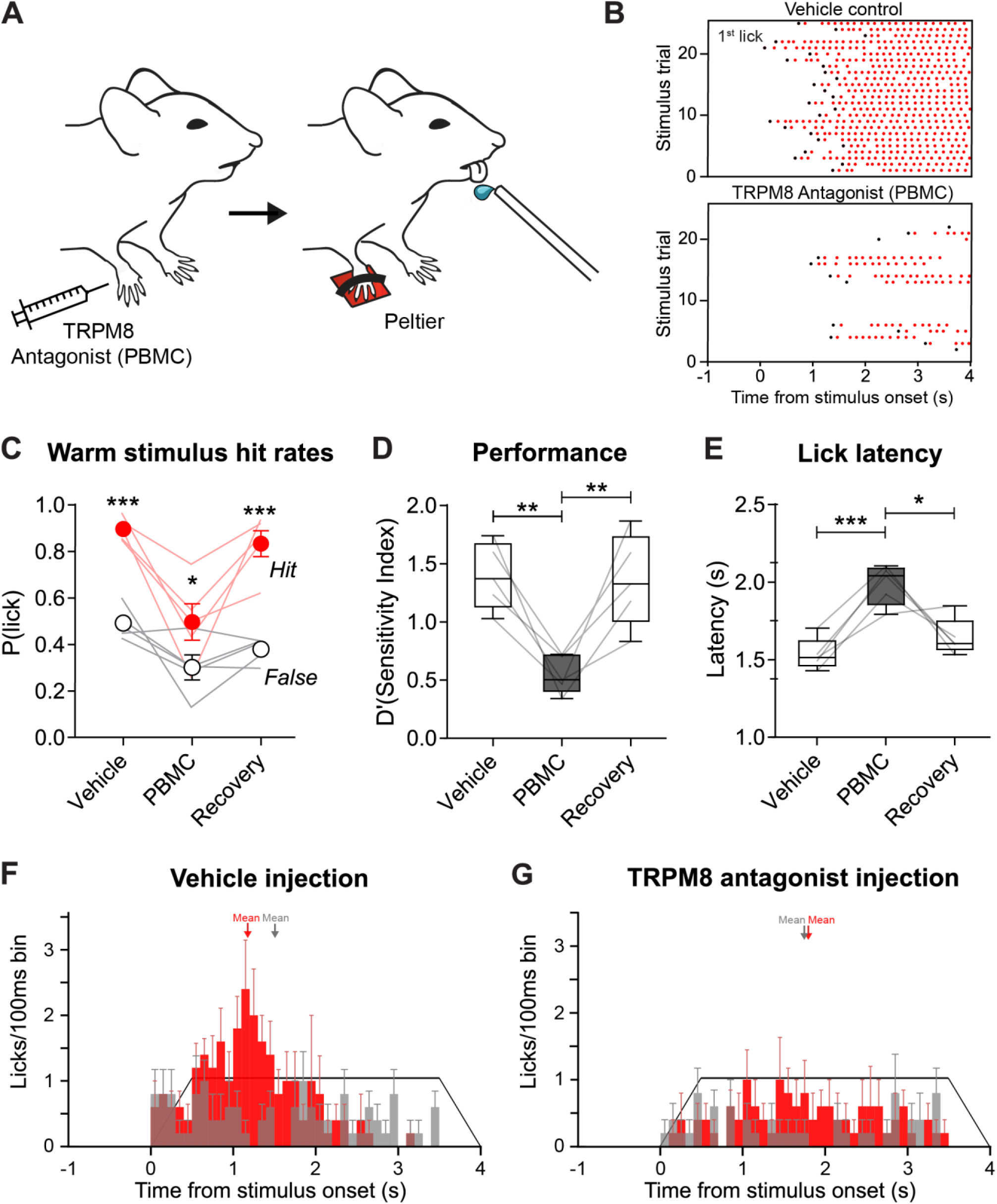
Forepaw administration of TRPM8 antagonist PBMC impairs warming perception. (A) The TRPM8 antagonist PBMC or DMSO vehicle control were injected into the forepaw of warming-trained wild type mice and afterwards their warming perception was assessed with the warming detection task. (B) Raster plots from DMSO vehicle-injected (top) and PBMC-injected (bottom) representative mice show that mice injected with a TRPM8 antagonist missed many of the trials. (C) Hit and false alarm rate differences in DMSO- and PBMC-injected mice were statistically significant, although the low hit rate observed in the PBMC-injected mice suggested a performance deficit. 24 hours later (Recovery), mice showed similar hit and false alarm rates to those of DMSO-injected conditions (n=5, two-way Anova with Bonferroni post-hoc analysis). (D) Sensitivity (D’) indices revealed an impaired performance in PBMC-injected mice, in comparison to the DMSO control group. Mice recovered warming perception one day later (n=5, paired t tests between PBMC and DMSO or Recovery groups). (E) In successful (hit) stimulus trials, PBMC-injected mice were slower to report the stimulus than in the DMSO-injected conditions. Similar to the performance, latencies decreased one day later (n=5, paired t tests between PBMC and DMSO or Recovery groups). (F) Average lick latency PSTH of DMSO vehicle-injected mice shows differences in the distributions between hits and false alarms, with a clear response peak only in the stimulus trials (red), and not in the catch trials (grey). (G) Population PSTH of PBMC-injected mice shows a much less clear response peak in the stimulus trials, and more similar hit (red) and false alarm (grey) latency distributions, further indicating a warmth perception deficit. *P < 0.05, **P < 0.01, ***P < 0.001. Data = mean ± SEM

## Discussion

While the afferent pathways and ion channels necessary for cooling perception have been studied extensively (Dhaka et al., 2007, 2008; Knowlton et al., 2013; McKemy et al., 2002; Milenkovic et al., 2014), far less is known about the molecular and neuronal pathways mediating innocuous warmth perception (Bokiniec et al., 2018; Filingeri, 2016). Here we show that mice learn to report warming stimuli delivered to the paw with similar perceptual thresholds to humans, and easily discriminate forelimb skin warming from cooling. We recorded the responses of afferent neurons innervating the forepaw with perceptually relevant thermal stimuli and characterized the classes of neurons that convey information on warming. We found two classes of sensory neurons that signal warming: a sub-population of polymodal C-MH fibers and most of the C-MHC fibers. No dedicated thermoreceptors were found to be activated by warming in mouse forepaw skin. Almost all warm sensitive C-MHC fibers also signal perceptually relevant cooling, thus these neurons provide an ambiguous signal that can only indicate that temperature has changed. Parallel information from warm sensitive C-MHs or cool sensitive C-MCs could then signal the direction of the thermal change. We also examined the impact of deleting *Trpm2 and Trpv1* genes on the sensory coding of perceptually-relevant skin warming (Tan and McNaughton, 2016; Yarmolinsky et al., 2016). Loss of TRPM2 or TRPV1 did not significantly impact the coding of warm by C-MH or C-MHC neurons, and *Trpv1* knockout mice show normal warm perception, whilst *Trpm2* knockout mice show mild warm perception deficits. However, there was a complete absence of warmth perception in mice lacking the cool-sensitive ion channel TRPM8: an effect that could be reproduced by acute pharmacological block of TRPM8 channels. The loss of TRPM8 channels diminished the coding of perceptually relevant cooling by both C-MC and C-MHC afferents. Together, our data indicate that warmth and cooling perception requires co-activation of an ambiguous low threshold channel (C-MHC) with polymodal C-fibers tuned either to skin warming (C-MH) or cooling (C-MC). Our data cannot be reconciled with labelled lines for warm and cool perception.

### Non-painful warm and cool perception is similar in mouse and human

Using a thermal training task we have shown that mice have remarkably similar warm and cool perceptual abilities to humans. Mice can detect forepaw skin warming of 1°C and skin cooling of 0.5°C from a 32°C baseline, values that closely match forearm thermal thresholds in humans (Stevens and Choo, 1998). Like in humans, the ability of mice to report forepaw warming is strongly dependent on spatial summation (Figure 1) (Filingeri, 2016; Stevens and Choo, 1998; Stevens et al., 1974), and mice easily discriminate non-noxious warming from cooling stimuli. Mice show higher sensitivity to cooling compared to warming as evidenced by lower absolute cooling thresholds and their ability to discriminate cooling applied to small skin areas with higher reliability than for warming (Figure 1). It has long been observed that in humans the perception of skin cooling is more acute and reliable than for warming (Stevens and Choo, 1998). The remarkable similarity in thermal perceptual ability between mice and humans strongly suggests that both afferent coding and the central processing of temperature in mice and humans have a common neural basis.

### Innocuous skin warming is signalled by two populations of polymodal C-fibers

Using warming as a search stimulus, we were able to identify the sensory afferents that convey perceptually relevant information to the brain. Around 20% of thermosensitive afferents in the hind paw and forepaw were classified as C-MHC neurons. Almost all C-MHC neurons signalled both cooling and warming and as such might signal that skin temperature has changed, because they show increases in firing rates to both warming and cooling, they are unlikely to provide information about the direction of temperature change. In addition, warming (defined here as 32-42°C) activated a sub-population (~70%) of C-MH fibers that can signal the direction of the temperature change. We did not record from any dedicated warming receptors with physiological properties similar to warm fibers recorded from the monkey hand and face (Darian-Smith et al., 1979b; Hallin et al., 1982; LaMotte and Campbell, 1978). It may be that dedicated warm-specific thermoreceptors do exist in the mouse skin, but are so rare that we have failed to sample them, however, even large scale imaging of thousands of DRG neurons to thermal stimuli have failed to convincingly identify neurons that respond only to warming (Chisholm et al., 2018; Wang et al., 2018; Yarmolinsky et al., 2016). In a now classic paper, LaMotte showed that sparse coding of warming by dedicated thermoreceptors in the monkey hand might account for psychophysical performance in humans (LaMotte and Campbell, 1978). However, warm receptors are also very rare in human skin, in one study just 5/125 C-fibers were found to exhibit the classic features of a dedicated warming receptor (Hallin et al 1981). In contrast, both polymodal C-MH and C-MHC fibers with physiological properties similar to those described here are very common (>40% of total C-fibers) in human skin (Campero and Bostock, 2010; Campero et al., 1996; Van Hees and Gybels, 1981). Information about skin warming must be spatially integrated in the spinal cord or brain as information carried in spike trains from single fibers are too sparse and unreliable to provide information about skin warming around the perceptual threshold of 1°C. Indeed, here we have calculated that the activation of more than 400 forepaw C-fibers by a 10°C warming stimulus may be required for the animal to reliably perceive warming. It is conceivable that one or more rare and specific thermoreceptor populations (dedicated cool and warm fibers) could instead provide the afferent drive for thermal perception, but so far no studies in rodents have revealed the existence of such afferents. A possible exception is a report of dedicated warm and cool sensitive C-fibers innervating the rat scrotum (Hellon et al., 1975), but thermosensation in this organ may not be specialized for perception.

### Heat activated TRP channels are not required for warm sensing

A recent study showed that genetic ablation of *Trpm2* abolishes thermal sensitivity in cultured DRG neurons in the range of 32-42°C *in vitro* (Tang and McNaughton, 2016). However, here we did not observe any loss in the sensitivity of C-MHC or C-MH fibers to innocuous warming or to noxious temperatures in the forepaw of *Trpm2^−/−^* mice (Figure 6). In contrast, we observed a reduction in the number of cold sensitive afferents in these mice, but the firing rates of the remaining cold sensitive fibers in were not different from those in controls (Figure 6). Similarly in *Trpv1^−/−^* mice we observed no impairment in the ability of C-MHC or C-MH fibers to code non-noxious temperatures, but we did observe a clear and specific impairment in the ability of C-MH fibers to code temperatures moving into the noxious range (>42°C) (Figure 6). This finding was robust and was also found for hind paw C-MH fibers (data not shown), but contrasts to recent reports that TRPV1 does not contribute to noxious heat sensing (Lawson et al., 2008; Woodbury et al., 2004). Interestingly, *Trpv1^−/−^* mice showed no impairment in their ability to report forepaw warming (Figure 5A,B) a finding at odds with conclusions made by Yarmolinsky et al (2016). The specific deficit in noxious heat sensitivity of C-MH fibers in the absence of TRPV1 is, however, in good agreement with the mild behavioural deficits in reacting to noxious heat observed in these animals (Caterina et al 2000). Our results also indicate that the same population of C-MH fibers can contribute both to warming perception and noxious heat sensation. Thus the context in which C-MH activity occurs may determine whether heating stimuli are perceived as innocuous or painful.

### Cooling sensitive afferents are required for warm perception

In contrast to *Trpv1^−/−^* mice we show that the absence of TRPM2 was associated with a small but significant reduction in behavioral sensitivity to warming. This behavioral deficit was seen in the absence of changes in the sensitivity of afferents to warming. However, in *Trpm2^−/−^* mice there was a significant reduction in the incidence of cold sensitive polymodal afferents (C-MC (Figure 6A). Cooling-sensitive C-fibers are known to be predominantly TRPM8+ (Bautista et al., 2007; Dhaka et al., 2008), and we confirmed here that in the absence of *Trpm8* many fewer C-fibers were found that responded to cooling in the 32–22 degree range (Bautista et al., 2007; Milenkovic et al., 2014). Unexpectedly, we observed a complete lack of warmth perception both in TRPM8*^−/−^* mice and in WT mice following an acute inhibition for the TRPM8 channel, but no clear change in the warm responses of afferent fibers recorded in *Trpm8^−/−^* mice. These data suggest that the processing of warmth and the resultant percept may require a comparison or integration of information from cooling sensitive polymodal C-fibers with information from warm sensitive C-fibers. Interestingly, the severity of behavioral deficits in warmth perception correlates well with the degree of loss in cooling sensitive fibers in *Trpm2^−/−^* and *Trpm8^−/−^* mice. One model that may explain these results is that a comparator circuit driven by ambiguous input from C-MHC fibers together with either warming information (C-MH) or cooling information (C-MC) determines the nature, sensitivity and magnitude of the percept. In this model, the activation of the C-MHC population by innocuous temperature changes plays a critical role as this population of afferents may be the only one that can signal that the stimulus is both innocuous and temperature specific. Loss of cooling sensitivity in this C-MHC population which is most prominent in *Trpm8^−/−^* mice could lead to ambiguity in the processing of temperature signals by comparator circuits in the spinal cord or brain. Indeed, such a model is reminiscent of warm/cold comparator circuit recently identified in the fruit fly *Drosophila melanogaster* (Liu et al., 2015).

In summary, our study reveals that warming and cooling perception in mice and humans share many features. Our data show that activity in overlapping populations of cutaneous polymodal C-fibers is sufficient to drive non-noxious thermal sensations, without the need for labelled lines. Indeed, the strong correlation between warm and cooling perceptual performance in humans (Frenzel et al., 2012; Stevens and Choo, 1998) could also be explained by the existence of a comparator circuit that is driven by both cool and warm sensitive C-fibers. Future work can now focus on elucidating the location and the mode of operation of central circuits mediating innocuous temperature perception using information from afferents that may also signal pain.

## Acknowledgements

This work was supported by the European Research Council (ERC-2017-ADG-789128, G.R.L.; ERC-2015-CoG-682422, J.F.A.P.), the European Union (3×3Dimaging 323945, J.F.A.P.), the Deutsche Forschungsgemeinschaft (Exc 257 NeuroCure, G.R.L., J.F.A.P.; FOR 2143, J.F.A.P.), the Thyssen Foundation (J.F.A.P.) and the Helmholtz Society (G.R.L., J.F.A.P.) We would like to thank Bettina Purfürst for help with electron microscopy, Valerie Bégay, Janett König and Charlene Memler for assistance with mouse breeding, mouse import and training, and Florian Rau for help with programming.

## Author Contributions

Conceptualisation, G.R.L, R.P.M, J.F.A.P and F.S; methodology, G.R.L, R.P.M, A.U, J.F.A.P, F.S and J.W.; formal analyses, R.P.M, F.S, J.W, A.U and R.E; Software, R.P.M; investigation, R.P.M, F.S, J.W and R.E.; writing original draft preparation, G.R.L, R.P.M, J.F.A.P and F.S; funding, G.R.L and J.F.A.P; supervision, G.R.L and J.F.A.P.

## METHODS

As Lead Contacts, James Poulet / Gary Lewin will fulfill any requests for further information, resources or reagents. Please contact.

### Animals

All experiments were approved by the Berlin animal ethics committee and carried out in accordance with European animal welfare law. Adult Wild-type C57Bl6/J mice and transgenic mice were used. The following strains of transgenic mice were used: 1) *Trpv1^−/−^* mice on a mixed background, from Jackson Laboratories (B6.129×1-Trpv1^tm1Jul^) (Caterina et al., 2000). 2) *Trpm2^−/−^* mice on a mixed background (129/SvJ and C57Bl6/N), backcrossed with C57Bl6/J mice for several generations, kindly donated by Yasuo Mori, Kyoto University (Yamamoto et al., 2008). 3) *Trpm8^−/−^* mice on a mixed background, from Jackson Laboratories (B6.129P2-Trpm8^tm1Jul^) (Bautista et al., 2007). All mice were maintained on a 12h light/ 12h dark cycle.

### Head implanting of mice for behavioural training

Mice were anesthetized with isoflurane (1.5–2% in O_2_) and injected subcutaneously with Metamizol (200 mg per kg of body weight). Temperature of mice was monitored with a rectal probe and kept at 37°C using with a heating pad. A light metal support was implanted onto the skull with glue (UHU dent) and dental cement (Paladur). Mice were then placed in their home cage with Metamizol (200 mg/ml) in the drinking supply 1-3 days.

### Behavioral training

Initially, head implanted mice were habituated to head-restraint in the behavioral setup for three days with increasing restriction times (15, 30 and 60 minutes). During the second and third habituation sessions, the right forepaw was fixed to the ground with medical (cloth) tape, in order to habituate the mice to paw-restraint.

Next, mice were water restricted and they underwent two “pairing” sessions in consecutive days. In these, water rewards were given from a water spout paired to presentation of the thermal stimulus in the forepaw (via an 3×3 or 8×8 mm Peltier element stimulator); to build an association between stimulus and reward. Each session lasted 1 hour approximately.

Mice that had undergone habituation and pairing started behavioral training. During training, mice only got a water reward (4-7 μl) from the spout when they licked it during a timeout upon start of the stimulus (3.5 seconds). Catch trials (where no stimulus is presented but licks are counted as false alarms) were included, interleaved, as 50% of the total trials.

Performance was assessed by counting hits and false alarms. All trials were delivered at randomized time intervals between 3 and 30 s. A training session consisted of about 100 trials (50 stimulus + 50 catch). Baseline temperature was 32°C, and stimuli consisted on an initial ramp to reach goal temperature (0.5 seconds), a hold phase (3 seconds) and a phase in which temperature returned to baseline (0.5 seconds). it was increased or decreased in 10°C during stimuli. In threshold experiments, stimulus amplitude was reduced every day (e.g. 6, 4, 2, 1, 0.5°C).

For sound training of *Trpm8^−/−^* mice, a magnetic buzzer generated a sound stimulus of about 40 dB that lasted for as long as the thermal stimulus. In the mechanical stimulation training, a Piezo stimulator produced a 3.5 seconds long single contact with the glabrous skin of the forepaw, and mice were rewarded when they licked within a time window of the same length as the thermal training.

### Skin-nerve preparation and sensory afferent recordings

Cutaneous sensory fiber recordings were perfumed using the *ex vivo* skin nerve preparation. Mice were euthanized by CO_2_ inhalation for 2-4min followed by cervical dislocation. In experiments using *Trp* knockout mice and C57/Bl6J control mice, the saphenous nerve and shaved hairy skin of the hind limb were dissected free. In forepaw experiments, the forepaw glabrous skin and innervating medial and ulnar nerves were dissected in a separate group of C57/Bl6J control mice. Skin and nerve samples were placed in an organ bath of 32°C perfused with a synthetic interstitial fluid (SIF buffer): 123mM NaCl, 3.5mM KCl, 0.7mM MgSO_4_, 1.7mM NaH_2_PO_4_, 2.0mM CaCl_2_, 9.5 mM sodium gluconate, 5.5mM glucose, 7.5mM sucrose and 10mM HEPES (pH7.4). The saphenous/medial and ulnar nerves were placed in an adjacent chamber in mineral oil, where fine filaments were teased from the nerve and placed on the recording electrode.

The receptive fields of individual thermosensory units were identified by pipetting hot (~48°C) and cold (~5°C) SIF buffer onto the surface of the skin. Electrical stimuli (1Hz, square pulses of 50-500ms) were delivered to unit receptive fields to classify them as C-fibers (velocity <1.2m/s), A-delta fibers (1.2-10m/s) or A-beta fibers (>10m/s). To test mechanosensitivity of units, four 3 second duration ramp and hold mechanical stimuli of increasing amplitude (20-400mN) were delivered using a computer controlled nanomotor^®^ (Kleindieck, Germany).

Next, to test thermal responses of units, a computer controlled Peltier device with a 3×3mm contact point (custom device built by Yale School of Medicine Instrumentation Repair and Design) was placed on the centre of the unit receptive field and a series of thermal stimuli were applied. In hairy hindpaw skin experiments, a heat ramp from 32 to 48°C (1°C/second) and a cold ramp from 32 to 12°C (1°C/second) was used. Average responses were obtained from three heat and cold ramps, with 2 minute intervals between each stimuli. In forepaw experiments, thermosensory unit receptive fields were stimulated with heat ramps which matched behavioural experiments: 0.5s ramp, 3s hold, and 0.5s ramp to baseline. 32-42°C heat ramps and 32-22°C cold ramps were given, and if units responded to these stimuli then a series of warm and/or cool ramps were given which decreased the amplitude by 2°C (e.g., 32-40°C, 32-38°C etc), followed by 32-33°C and 32-32.5 heat ramps, and/or 32-31°C and 32-31.5°C cool ramps. Thermal ramps were repeated 3–7 times, depending on the recording, to create average cell responses. Sensory fiber receptive fields were also stimulated using 1°C/second 32-48°C heat and 32-12°C cold ramps. Cells which exhibited signs of wind up or spontaneous activity after multiple stimulations were discarded from analysis.

### Transdermal injections in the forepaw

Mice that had been head implanted and trained (6 sessions) to report non-painful thermal stimuli in the forepaw were briefly anesthetized with isoflurane (1.5–2% in O_2_). Once the pain reflexes were absent due to the anaesthesia, 10 µL of solution were injected transdermally into the plantar side of the right forepaw, using a syringe of gauge 30G (0.3mm). Afterwards, mice recovered from anaesthesia. 15 minutes after the injection, all mice were active and were tested in the thermal perception task. As in all behavioural experiments described here, thermal stimuli were delivered to the right forepaw.

To control for the possible effects of the injection procedure and the anaesthesia, mice were injected in two occasions in different days: once with a solution in which the TRPM8 antagonist PBMC was absent (DMSO control); and once with a solution containing the drug (PBMC group). The injected solutions consisted of 4µL of DMSO with 0.1 mg of PBMC diluted in 6µL of saline (PBMC injection) and 4µL of DMSO in 6µL of saline (DMSO control).

### Electron microscopy

For electron microscopy images of the medial and ulnar nerves, animals were perfused and nerves was dissected and fixed in 4% PFA and 2.5% glutaraldehyde and contrasted with osmium tetroxide before embedding in Technovit 7100 resin (Heraeus Kulzer, Wehrheim, Germany). Ultra-thin sections captured at 5600x magnification. Myelinated nerve fibres and unmyelinated fibres were counted and measured using Fiji/ImageJ.

### Quantification and Statistical Analyses

#### Analysis of behavior

Licks were recorded with a sensor at the tip of the water reward spout. A thermocouple wire placed at the interface Peltier-forepaw skin measured the temperature during the training sessions. In stimulus trials, a hit was counted when there was a lick within the window of opportunity (3.5 seconds) after the start of the stimulus. During catch trials, a false alarm took place when there was a lick during an equally long window of opportunity.

To assess whether mice successfully learnt the detection task, hit rates were compared to false alarm rates within the same training session. Latencies to respond to stimuli were quantified and compared between groups as an additional measure.

To quantify performance in the detection tasks, we used D’ (sensitivity index) instead of the percentage of correct trials, in order to take into account bias in the licking criterion (Carandini and Churchland, 2013). To calculate d’, the following formula was used: D’ = z(h) – z(fa), where z(h) and z(fa) are the normal inverse of the cumulative distribution function of the hit and false alarm rates, respectively. To avoid infinity d’ values, when all trials were reported (rate = 1) or none of them was (rate = 0), the rates were replaced by (1–1/2N) or (1/2N), respectively, where N is the number of trials the stimulus was presented (Macmillan and Kaplan, 1985).

The z scores for hit and false alarm rates were calculated with OpenOffice Calc (Apache Software Foundation) using the function NORMINV.

Behavioral data was collected used custom-written routines in Lab View at 1 kHz sampling rate, and custom-written Python scripts were used for analysis.

#### Analysis of skin-nerve recordings

Cutaneous forepaw and hindpaw thermosensory units were categorized based on their conduction velocity and responses to thermal and mechanical stimuli.

Single unit recording thermal data points represent a mean response of >3 stimuli. Thermal and mechanical thresholds of units were calculated as the temperature or mechanical amplitude required to elicit the first action potential. In forepaw experiments, heat and cold-evoked firing activity was compared between different fiber populations (e.g., C-mechanoheat (C-MH) versus C-mechanoheatcold (C-MHC)). In hindpaw experiments, population responses of units recorded from wild type control and *Trp* knockout mice were statistically compared. Spike histogram graphs represent pooled data from multiple responses within and between C-fiber recordings in different animals.

#### Quantification of nerve axon counts and paw fiber density

Myelinated and non-myelinated axons were counted in 15 sections of nerves from each of 4 animals. To estimate the total number of axons per nerve, the cross sectional area of the nerve was measured and axon count data was extrapolated based on the area of the nerve quantified. The area of the glabrous skin of the forepaw and the hindpaw were measured, and fiber density was calculated from the axon number divided by the paw region area. This measurement is an over-estimation as it does not exclude proprioceptive fibers and somatosensory fibers which innervate deeper tissues such as joints and muscle. Proportions of thermoreceptive fibers were taken from skin nerve-preparation recordings and included into the calculations.

#### Statistical tests

Statistical analyses were performed with GraphPad Prism 5.0/6.0 and Python. Statistical tests for significance are stated in the text, and include two-way repeated measures ANOVA with Bonferroni’s post hoc test, Student t test, Mann Whitney test and Wilcoxon matched pairs test. Kolmogorov-Smirnov test was used to assess normality of the data. Asterisks in figures indicate statistical significance: *p<0.05, **p<0.01, ***p<0.001.

