## Supplementary Figures for "The sensory coding of warm perception"

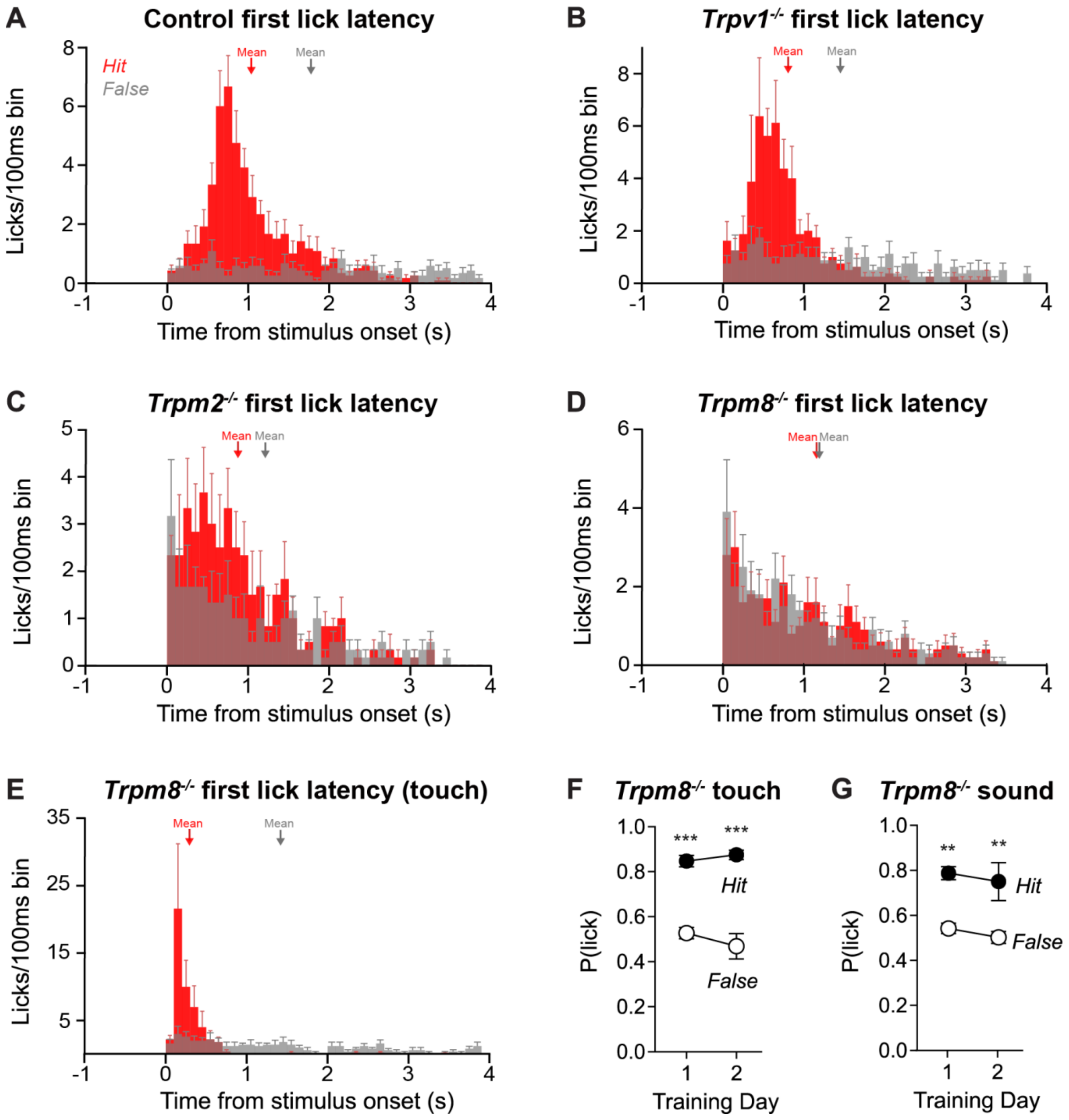


**Supplementary Figure 1. Warm detection latencies in control and TRP mutant mice.**

(A) Average PSTH of WT mice response latencies at training day 10 (n=12). Latencies in stimulus trials are shown in red, and catch trials are depicted in grey. Arrows show the distribution means. A clear peak can be observed only in the stimulus trials (red distribution), while the false alarm distribution appears flat.

(B) Average PSTH of *Trpv1*$^{-/-}$ mice response latencies at training day 10 (n=8). Hit and false alarm distributions appear very similar to those of WT mice.

(C) Average PSTH of *Trpm2*$^{-/-}$ mice response latencies at training day 10 (n=6). Note that, in comparison to wild type mice, the false alarm distribution is more prominent in comparison with the hit distribution. Moreover, the hit distribution appears more scattered and the peak is less prominent.

(D) Average PSTH of *Trpm8*$^{-/-}$ mice response latencies at training day 10 (n=10). Note that there is an almost equal distribution for both the stimulus trials and the catch trials, and the probability of licking decreased gradually after the stimulus onset, as a consequence of the frequent and random licking of the mice. Since the condition for a stimulus (or catch) to occur is that there is no lick during the pre-stimulus 3 seconds window, there are higher chances of a lick to occur shortly after the onset of the stimulus, if licking is very frequent.

(E) Average PSTH of *Trpm8*$^{-/-}$ mice response latencies at touch training day 2. Note that licks during catch trials are highly reduced, as opposed to hits, which show a fast and prominent peak (n=5). Interestingly, the responses to mechanical stimuli are much quicker than to thermal stimuli (shown in Fig. 2H).

(F) As opposed to warming stimuli, *Trpm8*$^{-/-}$ mice could learn to report mechanical stimuli in a detection task, as shown by high hit rates and comparatively low false alarms since the first training session (n=5, two-way repeated measures ANOVA with Bonferroni post-hoc tests).

(G) *Trpm8*$^{-/-}$ mice were also able to learn to report a sound cue in a detection task (n=5, two-way repeated measures ANOVA with Bonferroni post-hoc tests.)

Asterisks represent statistical significance between hit and false alarm rates, *P < 0.05, **P < 0.01, ***P < 0.001.

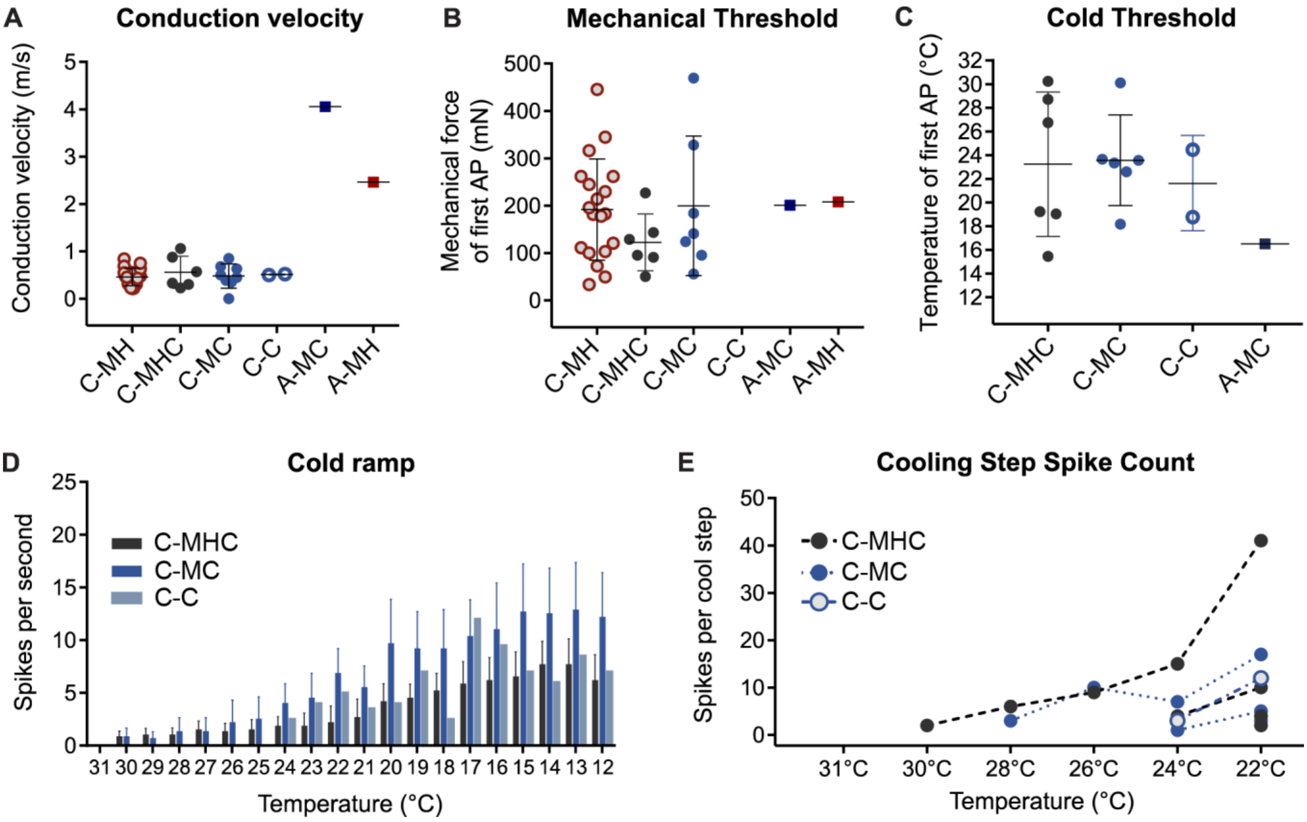


**Supplementary Figure 2. Cutaneous forepaw thermoreceptor responses to cool and mechanical stimuli.**

(A) Conduction velocities of thermosensitive afferent neurons innervating the mouse forepaw. C-MH n=19; C-MHC n=6; C-MC n=6; C-C n=2; A-MH n=1; A-MC n=1 (total n=35 from 10 animals).

(B) Mechanical thresholds did not differ between mechanoreceptive fiber populations. The only mechanically insensitive thermal units found were C-C units.

(C) Cold thresholds did not significantly differ between cold-responsive cell populations.

(D) Cool-evoked firing activity of populations of C-MHC, C-MC and C-C fibers during the 1ºC/s 32-12ºC cooling ramp.

(E) Cool-evoked firing activity of individual cool-responsive units are shown in response to 4s duration cool ramps used in behaviour experiments.

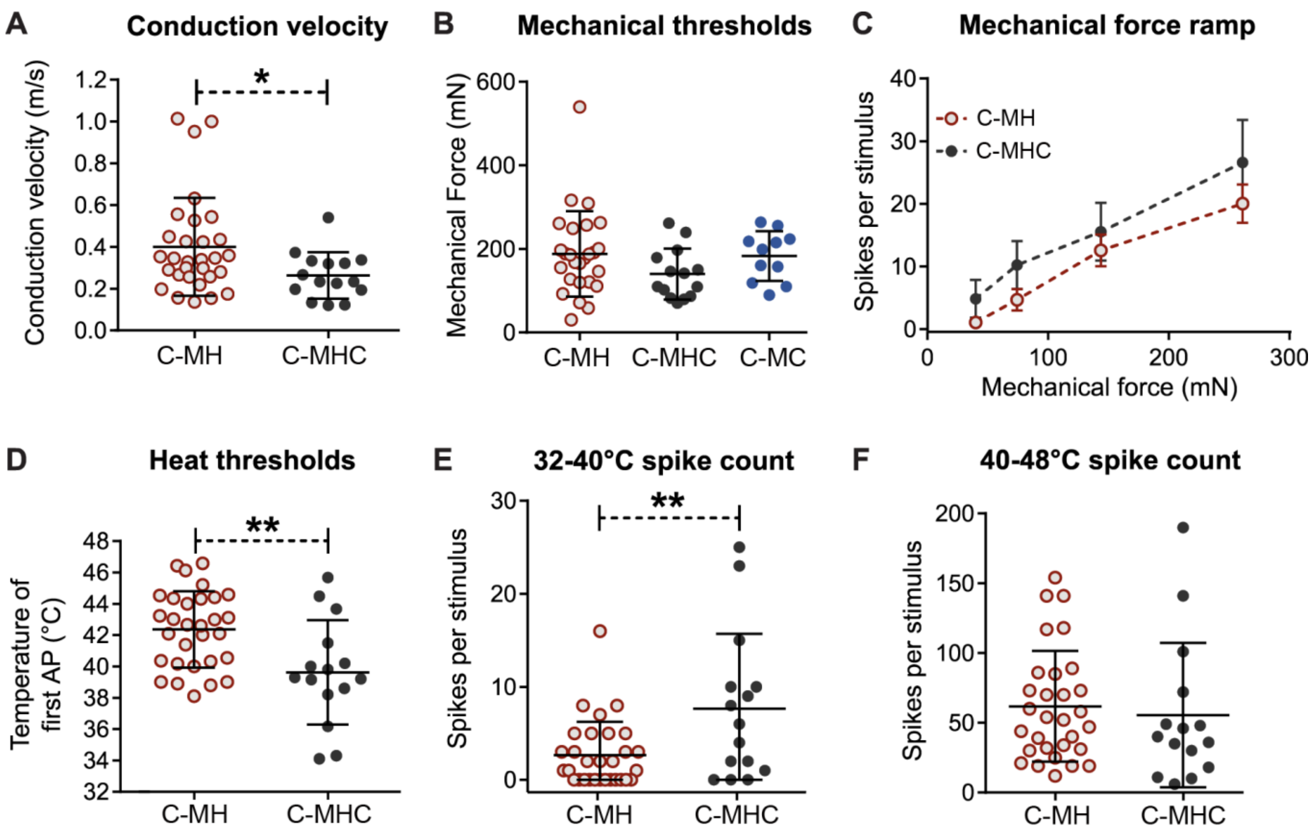


**Supplementary Figure 3. Cutaneous hindpaw thermoreceptive C-MHC fibers are more sensitive to warming than C-MH fibers.**

(A) Conduction velocities of saphenous thermosensitive afferent neurons innervating the hindpaw. C-MHC fibers were found to be significantly slower than C-MH fibers (n=30 C-MH and n=15 C-MHC fibers, unpaired t-test).

(B) Mechanical thresholds did not differ between C-MH, C-MHC and C-MC fibers innervating the hindpaw.

(C) Stimulus response curves during mechanical stimulation of increasing amplitude is shown for C-MH and C-MHC fibers.

(D) Heat thresholds were significantly lower in hindpaw-innervating C-MHC than in C-MH fibers using a 1$^{o}$C/s ramp between 32-48$^{o}$C (unpaired t-test).

(E) C-MHC firing activity was higher than C-MH activity during the warm 32-40$^{o}$C phase of the 1$^{o}$C/s 32-48$^{o}$C heating ramp (unpaired t-test).

(F) C-MHC and C-MH fiber responses to the noxious 40-48$^{o}$C range of the 1$^{o}$C/s 32-48$^{o}$C heating ramp were comparable.

Asterisks represent statistical significance between fiber types, *P < 0.05, **P < 0.01, ***P < 0.001.

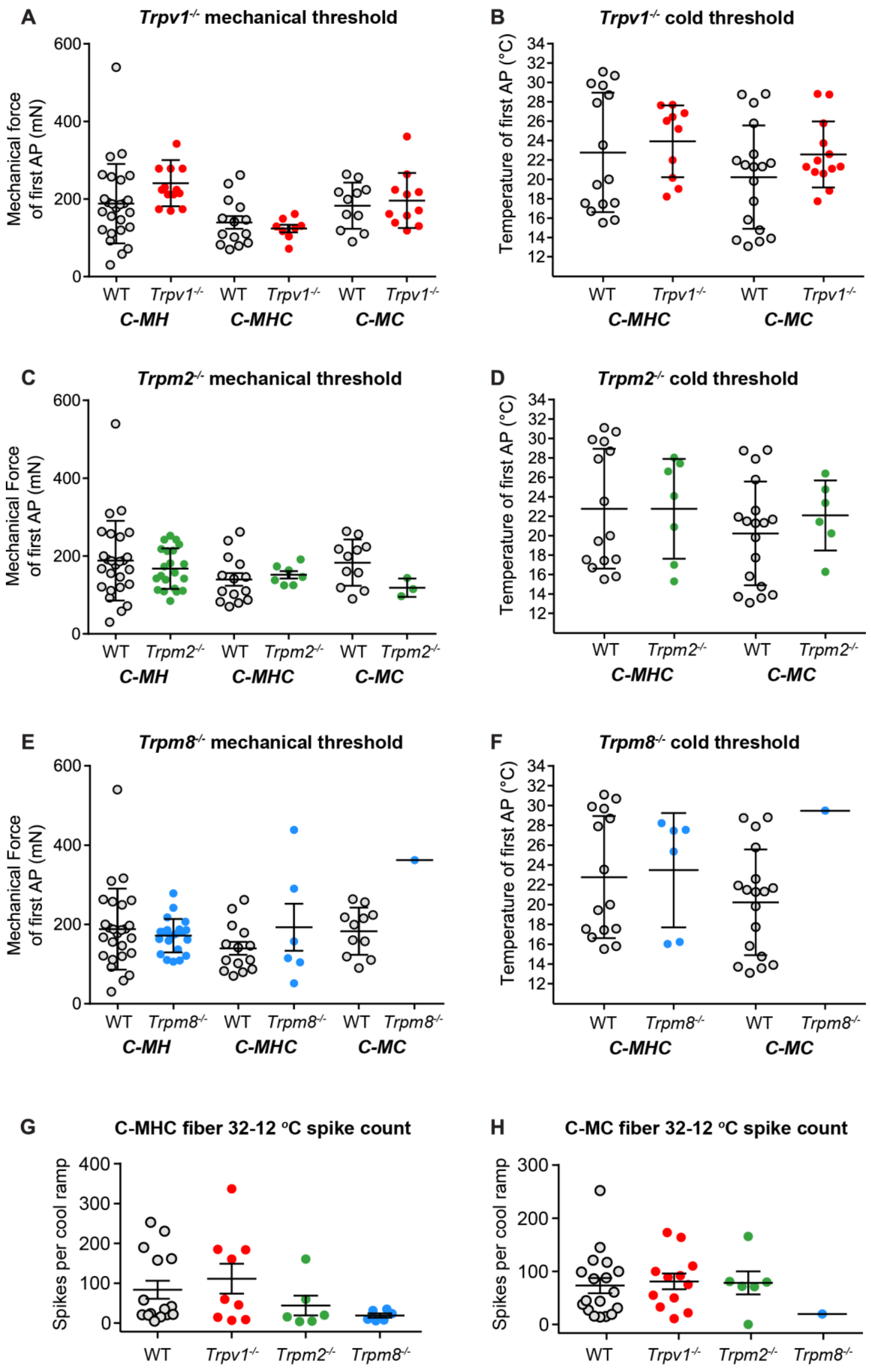
A
Trpv1-/- mechanical threshold
Mechanical force of first AP (mN)
600
400
200
0
WT
Trpv1-/-
C-MH
C-MHC
C-MC
B
Trpv1-/- cold threshold
Temperature of first AP (°C)
34
32
30
28
26
24
22
20
18
16
14
12
C
Trpm2-/- mechanical threshold
Mechanical Force of first AP (mN)
Trpm2-/-
D
Trpm2-/- cold threshold
E
Trpm8-/- mechanical threshold
Trpm8-/-
F
Trpm8-/- cold threshold
G
C-MHC fiber 32-12 °C spike count
Spikes per cool ramp
400
300
200
100
0
H
C-MC fiber 32-12 °C spike count

**Supplementary Figure 4. Mechanical and cold thresholds of hindpaw C-fibers in TRP mutant mice.**

(A) Mechanical thresholds of *Trpv1*$^{-/-}$ fibers (n=14 C-MH; n=10 C-MHC; and n=13 C-MC) compared to wild type control (n=30 C-MH; n=15 C-MHC; and n=18 C-MC fibers).

(B) Cold thresholds of *Trpv1*$^{-/-}$ C-MHC and C-MC fibers compared to wild type control.

(C) Mechanical thresholds of *Trpm2*$^{-/-}$ thermoreceptors (n=26 C-MH; n=7 C-MHC; and n=6 C-MC fibers).

(D) Cold thresholds of *Trpm2*$^{-/-}$ thermoreceptors did not differ compared to control.

(E) Mechanical thresholds of *Trpm8*$^{-/-}$ thermosensitive fibers (n=24 C-MH; n=6 C-MHC; and n=2 C-MC fibers) did not differ compared to control.

(F) Cold thresholds of *Trpm8*$^{-/-}$ thermoreceptors.

(G) Total spike activity of C-MHC fibers during the 32-12$^{o}$C cold ramp.

(H) Total spike activity of C-MC fibers during the 32-12$^{o}$C cold ramp.

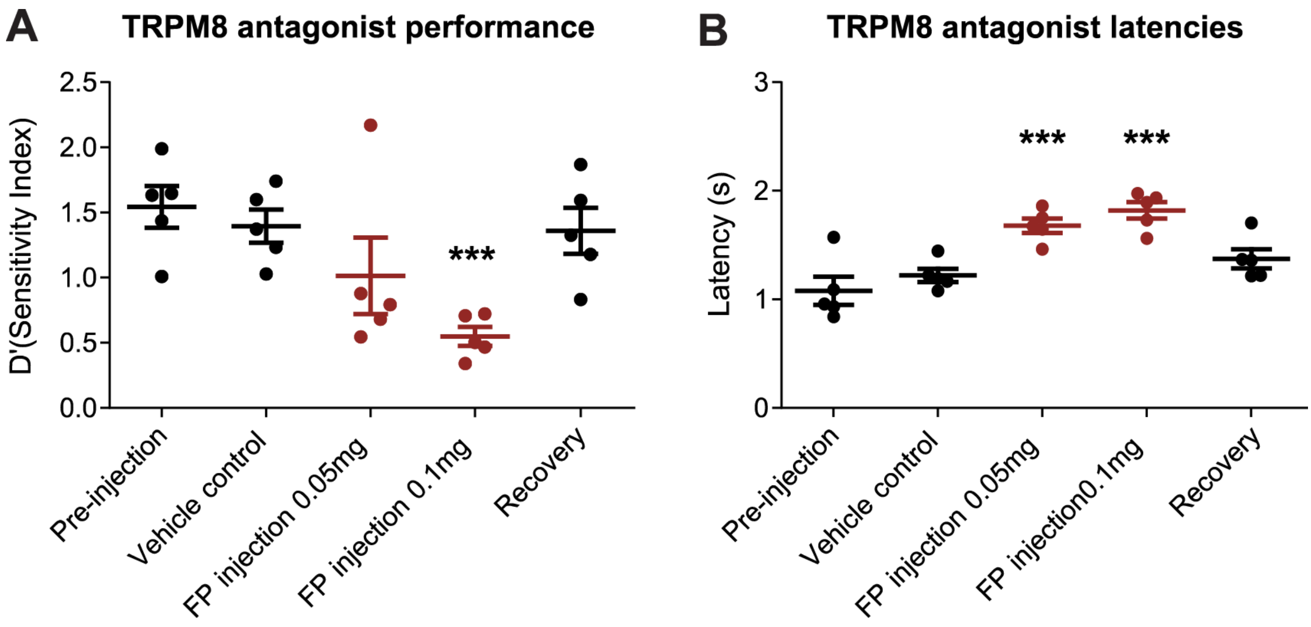


**Supplementary Figure 5. Performance of WT mice in a warming detection task following acute TRPM8 inactivation under different conditions.**

TRPM8 inactivation was tested via transdermal forepaw administration of the TRPM8 antagonist PBMC. Forepaw injections of 0.05 and 0.1 mg were tested, as well as injection of DMSO vehicle control. Pre-injection shows data from the last warming detection session prior to any treatment, and recovery depicts warming detection data from 24h after PBMC treatment.

(A) Sensitivity index (D') in the warming detection task was reduced in mice injected with 0.1mg of the TRPM8 antagonist PBMC in the forepaw, in comparison to when vehicle (DMSO) was injected transdermally in the forepaw (n=5 mice, paired t test).

(B) Mean latency to report warming stimuli was slower in mice injected with 0.1mg PBMC transdermally in the forepaw, in comparison to when DMSO vehicle solution was injected (n=5 mice, paired t test).

Asterisks represent statistical significance between vehicle control and TRPM8 antagonist injections, *P < 0.05, **P < 0.01, ***P < 0.001.
